# Residence-colonization trade-off and niche differentiation enable the coexistence of *Escherichia coli* phylogroups in healthy humans

**DOI:** 10.1101/2023.08.10.552787

**Authors:** Thibaut Morel-Journel, Sonja Lehtinen, Olivier Cotto, Rafika Amia, Sara Dion, Clarisse Figueroa, Jonathan Martinson, Pascal Ralaimazava, Olivier Clermont, Xavier Duval, Forough L. Nowrouzian, Seth Walk, Erick Denamur, François Blanquart

## Abstract

Despite extensive literature on the pathogenicity and virulence of the opportunistic pathogen *Escherichia coli*, much less is known about its ecological and evolutionary dynamics as a commensal in healthy hosts. Based on two detailed longitudinal datasets on the gut microbiota of healthy adult individuals followed over months to years in France and in the USA, we identified a robust trade-off between the ability to establish in a new host (colonization) and the ability to remain in the host (residence). Major *E. coli* lineages (phylogroups) exhibited similar fitness but a diversity of strategies, from strong colonizers residing for a few days in the gut, to poor colonizers residing for years. Strains with the largest number of extra-intestinal virulence associated genes and highest pathogenicity resided for longest in hosts. Moreover, the residence time of a strain was reduced more strongly when it competed with other strains of the same phylogroup than of different phylogroups, suggesting niche differentiation between them. To investigate the consequences of the trade-off and niche differentiation for coexistence between strains, we developed a discrete-state Markov model describing the dynamics of *E. coli* in a population of hosts. We found that the trade-off and niche differentiation acted together as equalizing and stabilizing mechanisms enabling the coexistence of phylogroups over extended periods of time. Our model predicted that a reduction in transmission (e.g. better hygiene) would not alter the balance between phylogroups, while disturbance of the microbiome (e.g. antibiotics) would hinder residents strains such as those of the extra-intestinal pathogenic phylogroup B2.3.

## Introduction

Understanding how bacteria diversify under the combined action of mutation and recombination is a long-standing issue in microbiology and evolutionary biology [1, 2]. Multiple mechanisms have been proposed to explain the observed diversity and the coexistence of bacterial strains that may exhibit fitness differences. Trade-offs between life-history traits can enable species to diversify into distinct strategies [3, 4]. Differentiation between ecological niches among bacteria can also favor diversity by allowing distinct strains to exploit the heterogeneity between hosts [5, 6]. For instance, the combination of a serotype-specific and non-specific host immune responses can generate host differences allowing coexistence among *Streptococcus pneumoniae* serotypes [7]. Other mechanisms not involving selection, such as a recombination rate negatively correlated with genetic distance, can lead to genetically distinct groups as well [1, 8].

The question of coexistence is especially critical for species such as *Escherichia coli*, which is not only a commensal bacteria of the human gut [9], but also an opportunistic pathogen. *E. coli* is responsible for various forms of diarrhea [10, 11] and extra-intestinal infections, including urinary tract [12, 13] and bloodstream infections [14, 15], causing around one million deaths worldwide each year [16–18]. The species is highly diverse and structured [19, 20], notably around lineages called ‘phylogroups’ [21, 22], which are largely monophyletic [9, 23] and differ in their virulence and antibiotic resistance profiles [24, 25]. Commensal and extraintestinal pathogenic *E. coli* are not strictly phylogenetically separated [26] and the transition to infection is the result of complex and hardly predictable interactions between several genetic, host and environmental factors [27]. Infections remain very rare events among the species, so that genes associated with pathogenicity (the probability to cause an infection) and virulence (the severity of infection) may evolve mainly as a result of their role in commensalism [28, 29]. Characterizing the dynamics of commensal *E. coli*, as well as their association with pathogenicity and virulence, is essential for understanding how the phylogenetic structure of *E. coli* emerged and for predicting the future evolution of the species.

This calls for precise characterization of the life-history traits of *E. coli*, which define the epidemiological dynamics in the human population – the rates at which strains colonize new hosts are cleared from them. These dynamics can be quantified through longitudinal follow-up of *E. coli* in healthy subjects, which unfortunately remains rare. Studies rather focus on diseased individuals [e.g. 30, 31], for example to assess the impact of antibiotic use on the microbiota [32, 33]. The initial colonization of infants during their first months of life has been studied [34–37], but longitudinal studies on healthy adults include a limited number of subjects [38–42].

The scarcity of longitudinal studies on healthy individuals can be explained by the complexity of their implementation. In addition to the need for a large number of subjects and of samples by subject, a single host can harbor several strains at the same time [43], multiplying the amount of work required to characterize each sample. In this study, we resolved this problem by using two complementary and remarkably detailed longitudinal datasets to characterize the dynamics of *E. coli* in healthy adults of high-income countries. The first one, previously analyzed by Martinson et al. [44], includes eight subjects followed over months to years in the USA with an extremely thorough characterization of the different clones present at the same time in each host. The second dataset, original to this study, includes 46 subjects followed over four to five months in France. Based on these datasets, we aimed to (i) quantitatively characterize the life-history traits of *E. coli* phylogroups, (ii) correlate these commensal traits with those associated with pathogenicity and virulence in the species and (iii) derive the consequences of our results for the epidemiological dynamics and potential coexistence of these phylogroups.

## Results

### Survival analyses

The first dataset we used to study the turnover dynamics of commensal *E. coli* (called thereafter the ‘French dataset’) included 415 clones sampled from 46 hosts over periods of 111 to 135 days. The second one (called thereafter the ‘USA dataset’) included 117 clones sampled from 8 hosts over periods of 245 to 849 days. We used the Clermont method [21, 22] to assign the clones of both datasets to phylogroups and additional PCR-based DNA fingerprinting techniques to differentiate clones of a same phylogroup within a single host (see Material and Methods).

We used survival models, specifically parametric accelerated failure time (AFT) models, to assess the impact of various factors on the rate at which clones colonized new hosts and their residence time once in their host. We performed one analysis per dataset (French or USA) and per response variable (colonization rate or residence time). For each analysis, we used the Akaike’s information criterion [AIC, 45] to select the combination of explanatory factors best explaining the data (see Supporting information). The explanatory factors potentially included in the best model were the following: the phylogroup of the clone considered, its host, the total number of other clones in the host, the number of other clones of the same phylogroup in the host and the total cell density in the host. We included the cell density of the clone considered in the analysis of residence time in the USA data only, as no precise estimate was available for the French data owing to the smaller number of colonies typed per sample.

### Determinants of the residence time

For both datasets, AFT models showed differences in clone residence times according to host and phylogroup (see Table S2 for the French dataset and Table S4 for the USA dataset). They also provided support for greater niche overlap between clones of the same phylogroup than between those of different phylogroups. Indeed, the residence time of a clone was significantly reduced by the presence of other clones in the host, but even more so if these other clones were of the same phylogroup (Table 1). According to the analysis of the French dataset, the residence time was divided by 1.45 (95%CI = [1.219, 1.717]) for each additional clone of a different phylogroup, and by 2.09 (95%CI = [1.594, 2.751]) for each additional clone of the same phylogroup. According to the analysis of the USA dataset, the residence time was divided by 1.68 (95%CI = [1.266, 2.358]) for each additional clone of different phylogroup, and by 5.09 (95%CI = [1.800, 14.423]) for each additional clone of the same phylogroup. The analysis of the USA dataset also showed that the cell density of a clone increased its residence time (Table 1), which was multiplied by 1.32 (95%CI = [1.002, 1.499]) when the clone density was multiplied by 10. Neither of the two AFT models selected based on AIC included total cell density as an explanatory factor.

**Table 1:** Z-score and p-values of the coefficients corresponding to the continuous explanatory variables in the AFT models of residence and colonization selected for the analyses of the French and USA datasets. Positive value of z-score indicate longer residence times/lower colonization rates as the factor increases. See Supporting information for the tables including qualitative factors, i.e. host and phylogroup.

| Dataset | Response variable | Explanatory factor | z-score | p-value |
| --- | --- | --- | --- | --- |
| French | Residence time | Total number of clones | -4.218 | < 0.0001 |
| French | Residence time | Number of clones of the same phylogroup | -2.654 | 0.0079 |
| USA | Residence time | Total number of clones | -3.444 | 0.0006 |
| USA | Residence time | Number of clones of the same phylogroup | -2.135 | 0.0328 |
| USA | Residence time | Log<sub>10</sub> of cell density | 1.983 | 0.0473 |
| French | Colonization rate | Total number of clones | -9.493 | < 0.0001 |
| USA | Colonization rate | Total number of clones | -9.775 | < 0.0001 |

To confirm the robustness of these results to the statistical method used, we performed additional analyses using semi-parametric Cox models with the same explanatory factors as those used for the AFT models, which yielded consistent results (see Supporting information).

### Determinants of the colonization rate

While we assumed that residence time was impacted by within-host dynamics alone, we considered colonization to be the result of inter-host transmission. Therefore, we expected the colonization rate of a phylogroup to be affected by its own frequency in the host population as a whole. To account for this, we corrected the colonization rates observed by the frequencies of phylogroups in their respective countries, based on cross-sectional external data (see Supporting information). We then performed the survival analyses using these corrected data.

As for residence times, AFT models showed differences in colonization rates according to host and phylogroup for both datasets (see Table S7 for the French dataset and Table S9 for the USA dataset). The two models also showed that the colonization rate was increased by the presence of other clones in the host (Table 1). The colonization rate was predicted to be multiplied by 1.76 (95%CI = [1.562, 1.970]) for each additional clone according to the analysis of the French dataset, and by 2.89 (95%CI = [2.342, 3.589]) for each additional clone according to the analysis of the USA dataset. However, the impact of other clones already residing in the host on the colonization rate was phylogroup-independent. Indeed, neither of the two AFT models selected based on AIC included the number of clones of the same phylogroup. They also did not select cell density in the host as an explanatory factor.

Again, we performed additional analyses using semi-parametric Cox models with the same explanatory factors as those used for the AFT models, which confirmed the results of the AFT models (see Supporting information).

### Trade-off between colonization and residence

To investigate the relationship between residence and colonization at the phylogroup level, we computed the estimates and standard errors of colonization rate and residence time predicted by the AFT models for each phylogroup (Fig 1) and quantified the association between them. The two life-history traits were negatively associated, both in France (Spearman’s *ρ* = *−*0.615, parametric bootstraping 95%CI = [*−*0.769, *−*0.238], *p* = 0.0016) and in the USA (Spearman’s *ρ* = *−*0.783, parametric bootstraping 95%CI = [*−*0.917, *−*0.599], *p <* 0.0001). This suggested a negative trade-off between the colonization and residence abilities at the phylogroup level. In other words, phylogroups were distributed along an axis ranging from those with a fast turnover (strong colonizers with short residence times) to those with a slow turnover (poor colonizers with long residence times). The arrangement of phylogroups along this trade-off was broadly conserved in France and the USA, with slow-turnover phylogroups including B2.3, D and F; intermediate phylogroups including A.2 and B1, and fast-turnover phylogroups including B2.1 (and B2.2 in the USA only, G in France only).

**Figure 1.**
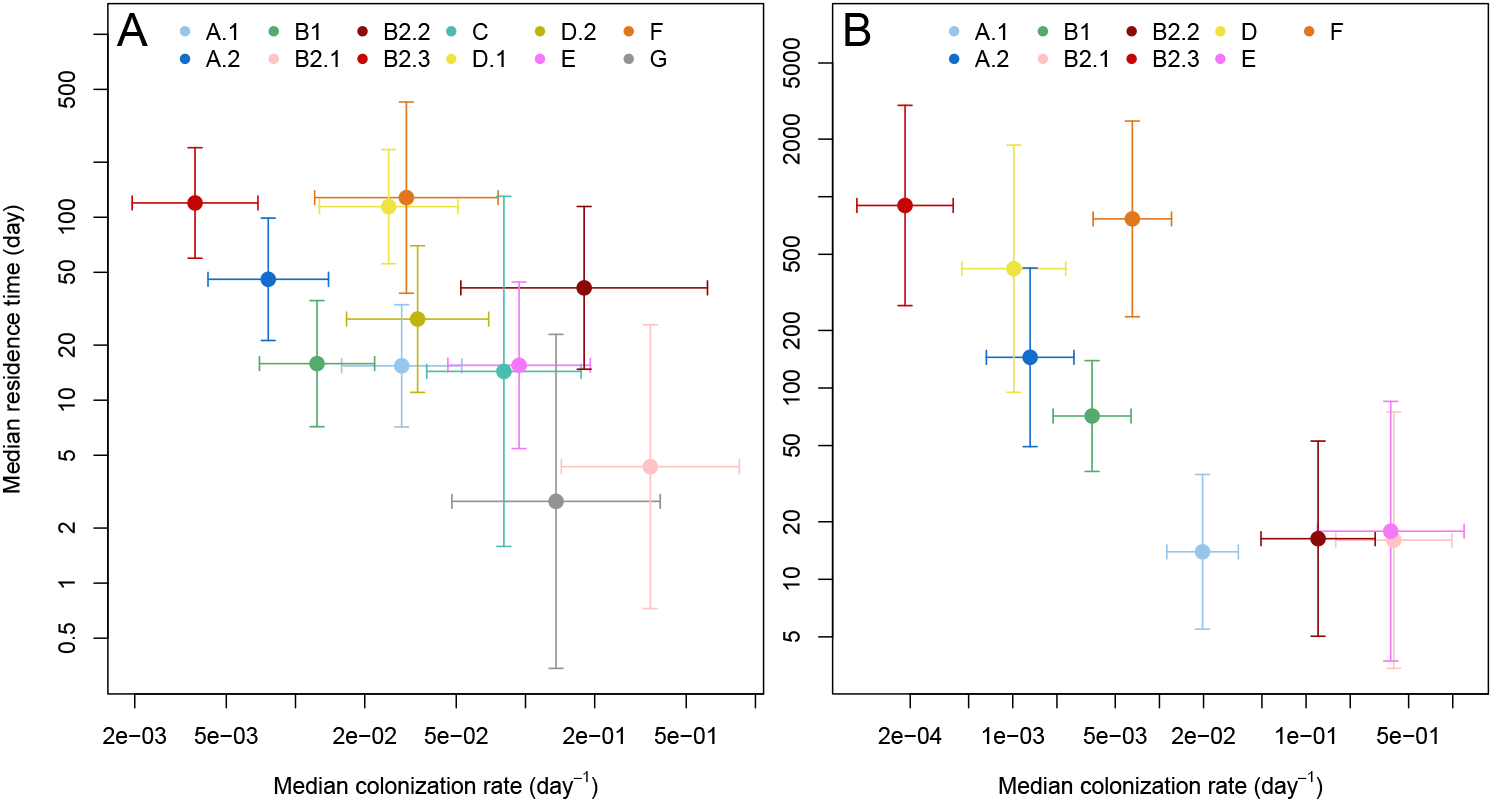
Median residence time as a function of the median colonization rate for each phylogroup appearing in the French (A) and the USA (B) datasets, for the average host and average values of the quantitative explanatory factors in their respective datasets. Estimates (dots) are represented with their 95% confidence interval (bars).

To further assess the robustness of the trade-off, we also analyzed colonization using colonization rates corrected by the frequencies of phylogroup in the datasets themselves rather than external data (see Supporting information). Using estimates from these models, we found similar negative correlations with the residence time, for the analyses of the French dataset (Spearman’s *ρ* = *−*0.769, parametric bootstraping 95%CI = [*−*0.832, *−*0.336], *p* = 0.0002) and the USA dataset (Spearman’s *ρ* = *−*0.733, parametric bootstraping 95%CI = [*−*0.900, *−*0.583], *p <* 0.0001).

Finally, we investigated whether the trade-off we identified in adults was also observed during the initial gut colonization of infants (see Supporting information). We analyzed an additional longitudinal dataset including infants sampled over a year after birth, previously analyzed by Ostblom et al. [36]. Results indicated that this trade-off was also visible in this specific context.

### Correlation between residence, pathogenicity and virulence-associated genes

The trade-off between colonization and residence has implications for the evolution of bacterial virulence, as the residents are also the most pathogenic bacteria, and tend to carry more virulence-associated genes (VAG) [46, 47]. We estimated pathogenicity (the odds ratio for the association between bloodstream infections and phylogroup) and the number of VAG carried for each phylogroup, based on a dataset previously analyzed by Marin et al. [48] and Burgaya et al. [49]. The estimated pathogenicity of phylogroups was positively correlated with residence for both datasets (French dataset: Spearman’s *ρ* = 0.518, parametric bootstraping 95%CI = [0.105, 0.741], *p* = 0.0131; USA dataset: Spearman’s *ρ* = 0.745, parametric bootstraping 95%CI = [0.217, 0.833], *p* = 0.0035)(Fig 2A, 2B). The total number of VAG carried was positively correlated with residence for the French data (Spearman’s *ρ* = 0.273, parametric bootstraping 95%CI = [0.049, 0.531], *p* = 0.0221) and the USA data suggested the same trend (Spearman’s *ρ* = 0.383, parametric bootstraping 95%CI = [*−*0.017, 0.533], *p* = 0.0527)(Fig 2C). Differences were observed in the strength of the correlation when considering genes associated with specific functions: those linked with protectins were the most strongly correlated with residence, while those linked with toxins were not. Genes linked with iron acquisition (*p* = 0.0473) and to adhesins (*p* = 0.0178) were significantly correlated with residence, for the French and USA datasets respectively.

**Figure 2.**
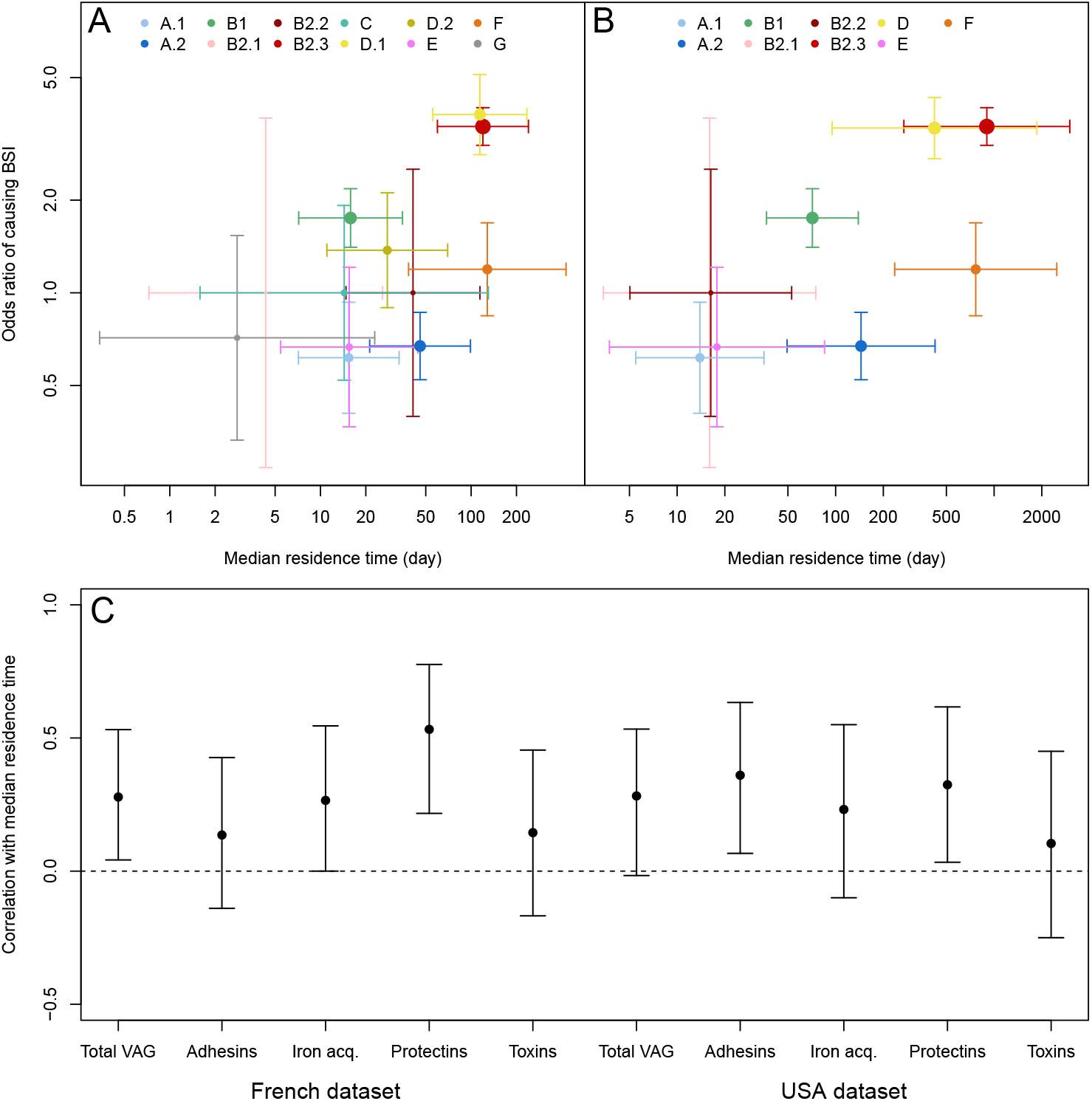
A and B: odds ratio of the association with bloodstream infections (BSI) as a function of median residence time for each phylogroup appearing in the French (A) and the USA (B) datasets. Estimates (dots) are represented with their 95% confidence interval (bars), with the size of the dots proportional to the number of samples of a given phylogroup used to estimate the odds ratios of causing BSI. C: correlation rate between the median residence time of each phylogroup estimated for the two datasets and its number of VAG (total or associated with adhesins, iron acquisition systems, protectins or toxins). Estimates (dots) are represented with their parametric bootstrap 95% confidence interval (bars). Correlations whose confidence interval does not include 0 are considered significant.

### Model of bacterial epidemiological dynamics

Following data analysis, we derived the implications of our results for the epidemiological and evolutionary dynamics of *E. coli*. We developed a continuous time, discrete-state Markov chain model describing the dynamics of clones in a host population and integrating our findings on colonization rates and residence times. In this model, discrete states were sets of the phylogroups of *E. coli* clones potentially present in the hosts. Phylogroups were numbered from 1 to *G* and states were numbered in increasing order of the number of clones in the host. For instance, an ‘empty’ host (without any clone) was in state *S*_1_ = {∅}, a host colonized by a single clone of phylogroup 1 in state *S*_2_ = {1}, a host colonized by a single clone of phylogroup 2 in state *S*_3_ = {2} and a host colonized by two clones of group 1 in state *S*_*G*+2_ = {1, 1}. The variable of the model was the probability *P*_*i*_(*t*) that a host was in state *S*_*i*_ at time *t*, which could also be interpreted as the proportion of an infinite population of hosts in state *S*_*i*_ at this time.

Transitions between states depended on clearance rates 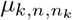 and colonization rates *λ*_*k*,*n*_(*t*):

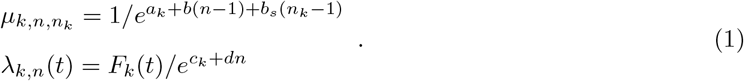

As inferred from the data, clearance was affected by three parameters: *a*_*k*_ was phylogroup-specific, *b* represented the impact of the number of other clones in the host *n −* 1 and *b*_*s*_ represented the impact of the number of other clones of the same phylogroup *n*_*k*_ *−* 1. In the French and USA datasets, we inferred negative values for *b* – clearance was faster the more clones were present in the host – and for *b*_*s*_ – clearance was faster the more other clones present belonged to the same phylogroup. The colonization rate of a clone was affected by the frequency of the phylogroup in the host population *F*_*k*_(*t*) and two parameters: *c*_*k*_ was phylogroup-specific and *d* represented the impact of the number of other clones in the host *n −* 1. In the French and USA datasets, we inferred negative values for *d*, meaning that colonization was easier when more clones were present in the host.

Each phylogroup was characterized by an intrinsic colonization-to-clearance ratio 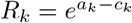, which was also its basic reproduction number. Indeed, when a single clone of phylogroup *k* was introduced in a large population of *H* empty hosts, the frequency of this clone was *F*_*k*_(*t*) = 1*/H*, the number of hosts to colonize was approximately *H* and the expected residence time of the clone was 1*/μ*_*k*,1,1_. Thus, the expected number of hosts colonized after this single introduction was:

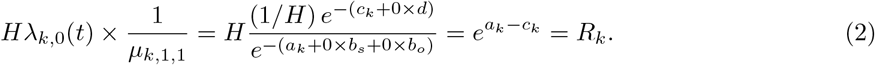

When competition between clones was the same irrespective of phylogroup (*b*_*s*_ = 0), the colonization-to-clearance ratio also predicted which phylogroup could invade others (see Supporting information). Indeed, the differences in colonization-to-clearance ratio *R*_*k*_ defined a ‘hierarchy’ between phylogroups, so that those with higher ratios could invade those with lower ratios. The existence of a trade-off then corresponded to several phylogroups with the same colonization-to-clearance ratio *R*_*k*_ but different strategies, e.g. from a fast turnover (high values of *a*_*k*_ and *c*_*k*_) to a slow turnover (low values of *a*_*k*_ and *c*_*k*_).

We explored with this mathematical model how this hierarchy between phylogroups was affected by non phylogroup-specific public hygiene measures (see Supporting information). On the one hand, the hierarchy was not affected by a measure resulting in a ‘multiplicative’ change, i.e. by multiplying the colonization or clearance rate by a constant factor. On the other hand, a measure adding a new source of clearance could alter the hierarchy of phylogroups by disproportionately affecting the phylogroup with a slower turnover. These changes in hierarchy could translate into variations in phylogroup frequencies. For instance, increasing the strength of this source of clearance eventually lead to the dominance of a fast-turnover phylogroup over a slow-turnover one (Fig 3).

**Figure 3.**
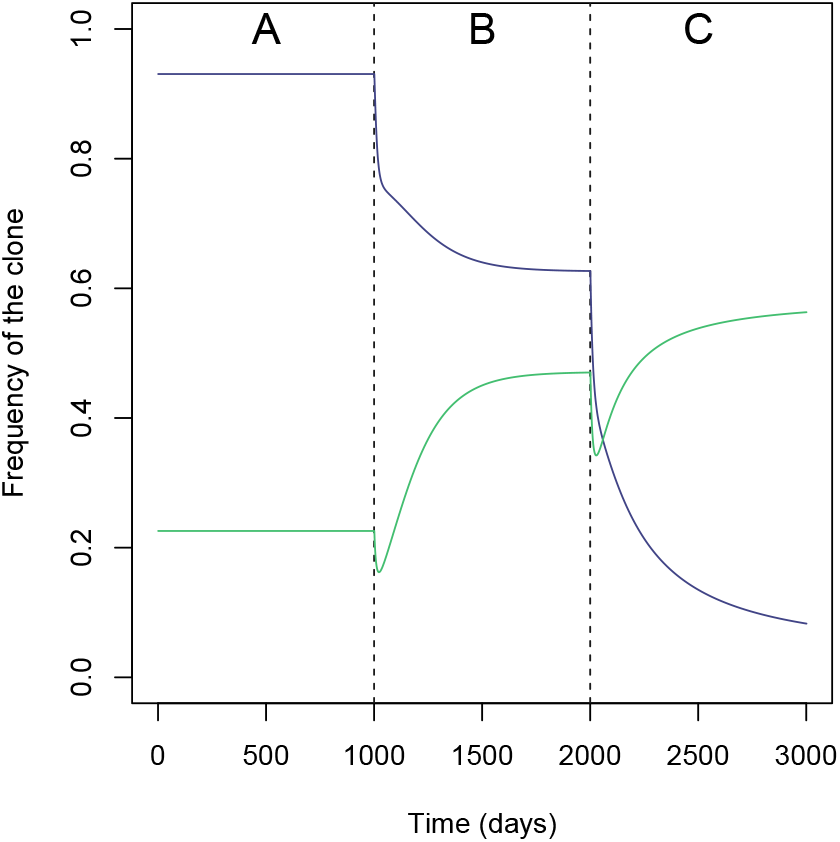
Variation in the frequencies of phylogroups *x* (purple) and *y* (green) over 3000 simulated days of the model with increasing levels of additive clearance (*θ*), with *R*_*x*_ = 3.5, *R*_*y*_ = 3, *b* = 0, *b*_*s*_ = *−*0.1 and *d* = 0. Three time-periods of 1000 days each (separated by dashed lines) are defined: no additive clearance (*θ* = 0) for *t∈* [0, 1000[(A), *θ* = 0.1 for *t* ∈ [1000, 2000[(B) and *θ* = 0.2 for *t* ∈ [2000, 3000[(C). Phylogroup *x* has a slower turnover than *y*. the phylogroup hierarchy is reversed for a high enough value of *θ*.

### Conditions for the coexistence of multiple phylogroups

We next assessed the conditions in which our model enabled coexistence between two phylogroups, depending on their respective colonization-to-clearance ratios. We quantified coexistence with two measures. Within-host coexistence *E*^*W*^ (*t*) ∈ [0, 1] corresponded to the proportions of hosts carrying clones of at least two phylogroups. Inter-host coexistence *E*^*I*^ (*t*) ∈ [0, ln (*G*)] was a Shannon index on the relative frequencies of the phylogroups in the host population:

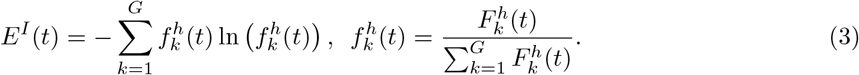

We considered a simple case of two phylogroups *i* and *j*, characterized by an average colonization-to-clearance ratio 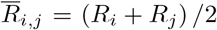 and a ratio difference *D*_*i*,*j*_ = *R*_*i*_ *− R*_*j*_. We performed discrete-time simulations of our model with a time-step of one day until an asymptotic state was reached. We assumed that this state to be reached when state probabilities essentially no longer varied, i.e. at time *t*\* such that 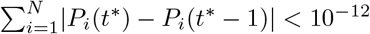. We computed the asymptotic within-host coexistence *E*^*W*^* and inter-host coexistence *E*^*I*^* at the steady state.

The results showed that only structurally unstable coexistence was possible if there was no additional competition between clones of the same phylogroup (*b*_*s*_ = 0, see Eq. 1), which was in line with the mathematical analysis of the model (see Supporting information). In that case, coexistence was maintained only if the two phylogroups had exactly the same colonization-to-clearance ratio, i.e. if *D*_*i*,*j*_ = 0. Stable coexistence was possible when there was additional competition between clones of the same phylogroup (*b*_*s*_ *<* 0). As *b*_*s*_ became more negative (greater niche differentiation), the range of values of *D*_*i*,*j*_ for which coexistence was stable widened, but within-host coexistence also declined (Fig 4A). Indeed, more negative values of *b*_*s*_ lead to higher average clearance rates overall, thus decreasing overall the number of clones per host, and the value of *E*^*W*^*. Inter-host coexistence was not impacted in the same way and *E*^*I*^* remained high as *b*_*s*_ became more negative (Fig 4B).

**Figure 4.**
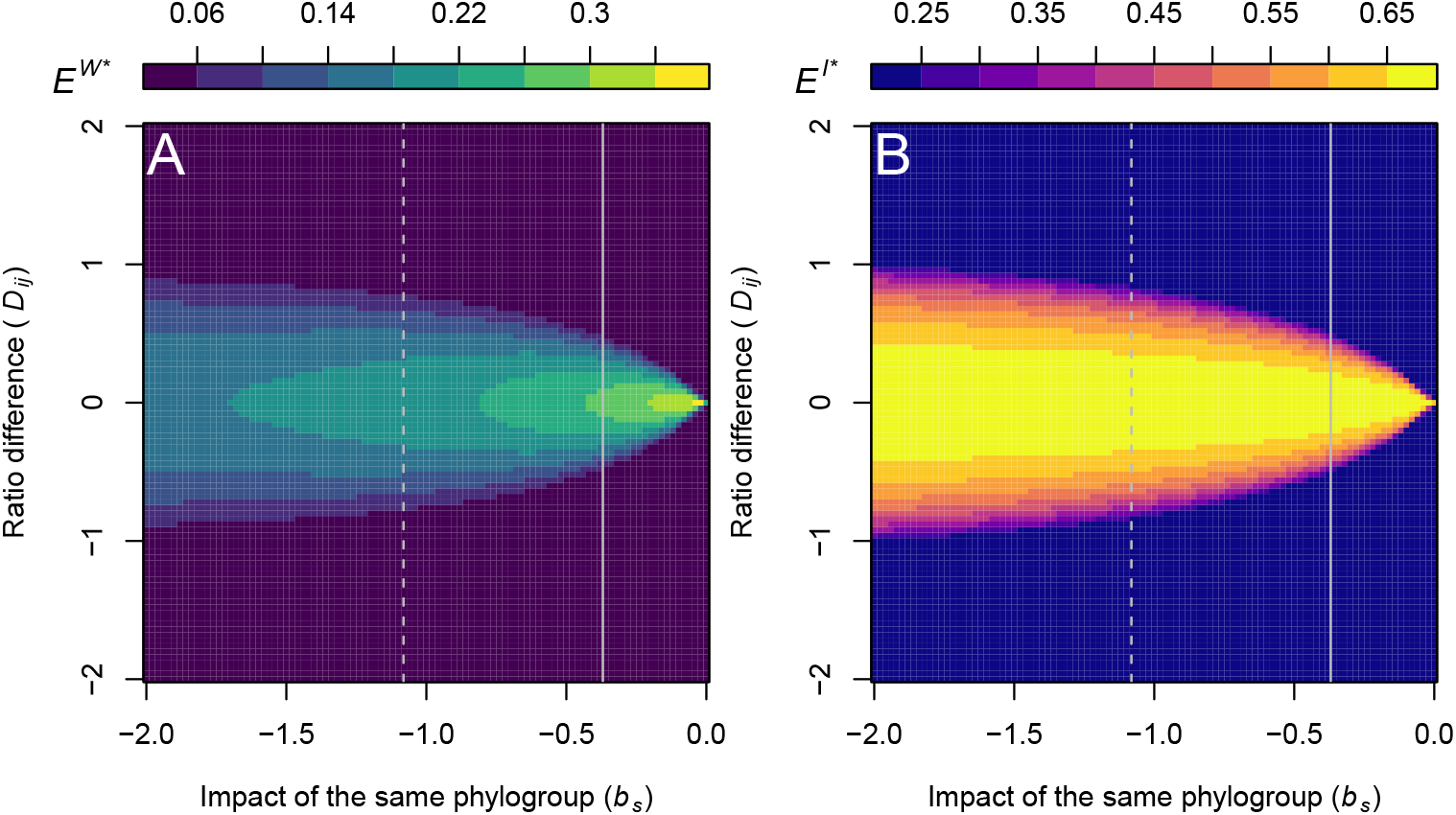
Asymptotic within-host coexistence *E*^*W*^* (A) and inter-host coexistence *E*^*I*^* (B) as a function of the ratio difference *D*_*i*,*j*_ and the strength of clones of the same phylogroup on clearance *b*_*s*_. Vertical lines correspond to the coefficients characterizing the impact of other clones of the same phylogroup from the best AFT models for the French (solid) and USA datasets (dashed). For these simulations, *R*_*i*,*j*_ = 2, *b* = 0, *d* = 0, *E*^*W*^* ∈]0, 1] and *E*^*I*^* ∈]0, ln (2)].

The simulation results presented in Fig 4 correspond to 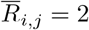, *b* = 0 and *d* = 0, but similar results were obtained for other parameter values. Higher values of 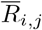 (the average colonization-to-extinction ratio), more positive values of *b* (lower clearance rate as the number of clones in the host increases) and more negative values of *d* (higher colonization rate as the number of clones in the host increases) all resulted in a higher overall number of clones per host, and therefore wider ranges of *D*_*i*,*j*_ for which coexistence was stable for a given value of *b*_*s*_. Lower values of 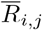, more negative values of *b* or more positive values of *d* had the opposite effect.

## Discussion

By analyzing longitudinal datasets on *E. coli* gut colonization dynamics in healthy adults from high-income countries, we identified two mechanisms explaining the stable coexistence of *E. coli* phylogroups. The first one is the trade-off between the colonization and residence abilities at the phylogroup level. Our results indicate that the ‘resident’ or ‘transient’ nature of *E. coli* clones, which has been observed for over 100 years [39, 50], is a property partly explained by phylogroups and inversely correlated with the ability to colonize hosts. This trade-off is equalizing *sensu* Chesson [51], meaning that it reduces differences in fitness between the phylogroups. In our model of colonization dynamics with human-to-human transmission, fitness is mathematically equal to the colonization-to-clearance ratio *R*_*k*_. Therefore, the coexistence of multiple phylogroups in a population at equilibrium implies that these phylogroups compensate shorter residence times with a higher colonization rate (or vice versa) to achieve similar fitness. Indeed, clones with lower fitness would not rise in frequency, while clones with larger fitness would invade and replace all other clones. This trade-off was also evidenced in infants followed during their first year of life, who have much simpler microbiome [52], suggesting that it does not depend on the presence of other species.

The second mechanism is niche differentiation, which acts as a stabilizing mechanism that can explain long-term coexistence [51]. In both datasets, residence is shorter when a clone is in a host together with another clone of the same phylogroup. This indicates stronger competition within phylogroups than between phylogroups, which could be explained by a greater overlap of ecological niches between clones that are phylogenetically closer. Interestingly, colonization does not exhibit the same dependence to the number of clones of the same phylogroup. This suggests that the niche overlap observed only affects clones after the colonization of a new host.

Our analyses also highlighted substantial heterogeneity between hosts in bacterial dynamics, consistent with the absence of ‘core resident microbiome’ initially reported by Martinson et al. [44] using the USA data. Clone turnover rates varied not only between hosts, but also over time within the same host. Both clearance and colonization were more frequent in hosts with a greater number of different clones. This was particularly apparent in the USA dataset, in which hosts experienced periods of stability with few clones, alternating with rapid successions of colonization and clearance events, during which the number of clones present temporarily increased. These periods of instability could be triggered by temporal variations in the suitability of the host for colonization or residence. They could also be triggered by the temporary destabilization of the microbial community due to the colonization of several clones in a short time span. These explanations are not exclusive and could both contribute to temporal variability. These transient periods of instability have also been observed over decades in a single host, with long periods of residence of clones, mainly of phylogroup B2, interrupted by rapid successions of clones, mainly of phylogroup B1 or A [42].

Although this study focuses on commensal *E. coli*, our results also bear implications for the evolution of pathogenicity. Previous studies have highlighted phylogroup B2, massively represented in our data by B2.3 (see Fig S1) as both resident and virulent [29, 53, 54]. We present a more general positive correlation between the residence time of a phylogroup, and both its pathogenicity and its number of virulence-associated genes. These results are consistent with the hypothesis that the virulence of *E. coli* as an opportunistic pathogen results from traits linked with its residence ability as a commensal [29, 55]. Our additional analyses focusing on genes associated with specific functions show that protectins, and to a lesser extent adhesins and iron acquisition systems, are positively correlated with residence, while the signal was unclear for toxins. Interestingly, while the most resident phylogroups were identified as the most likely to generate extra-intestinal infections, better colonizing phylogroups such as B1, A.1 or E have been associated with pathovars causing severe and acute diarrhea [56]. These pathovars may benefit more from the ability to colonize rather than from the ability to reside.

Our results might also have implications for the evolution of *E. coli* in response to public health measures. On the one hand, ‘multiplicative’ measures increasing overall clearance rate (or reducing overall transmission) rate by a given factor are not predicted to alter the fitness hierarchy between phylogroups. On the other hand, ‘additive’ measures creating new sources of clearance, such as antibiotics, should favor the best colonizers at the expense of the residents. This could explain the resilience of the microbiota in settings with high antibiotic use [57]. Conversely, a reduction in disturbances of the microbiota could explain an increase in the frequency of phylogroup B2, as observed in France [25, 48]. The disproportionate impact of antibiotics on the most resident phylogroups could favor less pathogenic ones, which are better colonizers than residents. It could also generate stronger selection for antibiotic resistance among residents [58]. Yet, we found no link between residence and the presence of resistance genes in our data [25, 48]. This differs, for example, from *S. pneumoniae*, another human commensal bacteria and opportunistic pathogen, where serotypes carried for longer are also more antibioresistant [58].

Our study has limitations. We focused on colonization and clearance rates, which likely result from many underlying mechanisms, such as resource uptake, bacterial warfare or predation by phages [59]. Moreover, residence as measured in our study corresponds to the ability for a clone to maintain at densities high enough to be detected in the dominant flora. The dynamics of subdominant clones remains unobserved and would be interesting to quantify in further studies. Increasing the number of clones characterized by sample is expected to improve the completeness of the diversity observed in the data, but at a cost, for instance on the number of hosts included. Our results are supported by congruent analyses on two complementary datasets, focusing respectively on a large number of hosts (French) or on a large number of clones characterized per sample (USA). To test whether sampling a limited number of clones could bias our results or limit power, we randomly sub-sampled the USA dataset to five clones per sample (see Supporting information). Neither the trade-off nor the niche differentiation were biased by sub-sampling. The trade-off was still robustly identified, with a consistent negative correlation between residence and colonization similar to the one observed with the complete data, while the statistical power to detect niche differentiation was reduced, presumably because of the reduced diversity in sub-sampled data.

Colonization encompasses all events between the shedding of a clone by a host and its detection at high enough densities in another host. In our model, this series of events is encapsulated in the colonization rate. This is a valid description if the time spent by bacterial cells in the environment between hosts is short. In this model, better survival in the environment would result in a higher colonization rate. However, our model cannot account for long residence time in the environment or non-human hosts, which are known to harbor distinct *E. coli* clones [60, 61] with dynamics potentially largely independent from the human hosts. We investigated how transmission of clones from an external environmental reservoir would affect the outcome of our model (see Supporting information). Results show that an external reservoir could stabilize coexistence by providing a potential additional source of rare strains in the population.

In conclusion, by analyzing longitudinal data on *E. coli* epidemiological dynamics in healthy adults from two separate datasets, we discovered general principles structuring the diversity of this species: major lineages align along a colonization-residence trade-off equalizing their fitness, and their differentiated niches in humans stabilize their long-term coexistence. We also discovered a strong correlation between residence and pathogenicity. Our results on high-income countries, where the phylogroup with the slowest turnover B2 is over-represented [9, 43, 62], would be nicely complemented by longitudinal analyses in other countries with radically different phylogroup frequencies. Finally, our work also suggests mechanisms by which the micro-evolutionary dynamics of *E. coli* can be altered, and opens perspective for the study of epidemiological and evolutionary dynamics of other human commensal bacteria and opportunistic pathogens.

## Material and Methods

### Longitudinal datasets

We used two longitudinal datasets collected from adults living in high-income countries to perform the survival analyses. The first one, referred to as the French dataset, was original to this study. See the Supporting information for further details about the data collection and Fig S9 for a graphical representation of the longitudinal data. This dataset included 415 clones collected from 46 healthy subjects in the Paris area over the course of 10 samples approximately every two weeks (14.9 days on average between samples, SD = 4.9 days) in 2021 and 2022. The clones collected were differentiated using multilocus variable number tandem repeat analysis (MLVA) [63]. Each clone was assigned to one of twelve phylogroups, following Clermont et al. [21, 22]: A.1, A.2, B1, B2.1, B2.2, B2.3, C, D.1, D.2, E, F and G.

The second dataset, referred to as the USA dataset, was previously analyzed by Martinson et al. [44]. See Fig S8 for a graphical representation of the longitudinal data. This dataset included 117 clones collected from eight healthy subjects over periods ranging from 245 to 849 days (11.7 days on average between two samples, SD = 11.1 days). The clones collected were differentiated using GTG5 repetitive element amplification [64]. Each clone was assigned to one of nine phylogroups following the same method as for the French dataset: A.1, A.2, B1, B2.1, B2.2, B2.3, D (which includes D.1 and D.2 as defined in the French dataset), E and F. Phylogroups C and G did not appear in this dataset, respectively because a single clone of group C was recorded and was discarded, and because group G was not defined separately in the original study.

To assess whether our conclusions hold for colonization dynamics in infants, we considered a third dataset previously analyzed by Ostblom et al. [36] and referred to as the infant dataset. See the Supporting information for details about this dataset and the analyses performed, and Fig S10 for a graphical representation of the longitudinal data. This dataset included 273 clones collected from 130 infants in Sweden at fixed times since birth (after 3 days, 1, 2 and 4 weeks, 2, 6 and 12 months). The clones were assigned to one of the four main phylogroups: A (including A.1 and A.2), B1, B2 (including B2.1, B2.2, B2.3) and D (including D.1 and D.2).

### Survival analyses

We used the software *R* [65] and the package *survival* [66] to perform survival analyses on the French and USA datasets. We used parametric accelerated failure time (AFT) models, which described the relationship between the time to an event – colonization or clearance – and a set of explanatory factors [67]. This type of model takes into account interval-censored data, i.e. the fact that colonization or clearance can occur between two samples. We counted the residence time of a clone (time to clearance) starting from its first appearance in the host. We counted the time to colonization from the first appearance of the last preceding colonizing clone. For clarity, the results are presented in terms of the colonization rate, defined as the inverse of this time to colonization. We corrected the times to colonization to account for the differences in colonization rate due to differences in phylogroup frequency in the general population, based on the frequencies of the phylogroup in France and in the USA. See the Supporting information for more details on the method. We excluded left-censored data, but included right-censored data (clones still present in the last sample of the host) for the AFT models of residence time.

We considered how colonization and clearance depended on two qualitative factors – the host and the phylogroup of the focal clone – as well as the following quantitative factors: the total cell density of *E. coli* (in CFU/g), the number of clones inhabiting the host (other than the focal clone), and the number of other clones of the same phylogroup (other than the focal clone). We also considered the cell density of the clone considered for the AFT models of residence time of the USA dataset, as this information was not available for the French dataset owing to the smaller number of colonies typed per time-point. Because of the wide variations in cell densities observed between samples, we used their log_10_-transformed values.

We performed model selection based on Akaike’s information criterion [AIC, 45] to identify the AFT model best explaining the residence times and times to colonization in the French and USA datasets. See the Supporting information for details about the model selection. In each case, we considered the models including any combinations of the explanatory factors presented above, with the time to the event following one of the three following distributions: exponential, Weibull and log-logistic. We ranked these models based on their ΔAIC, i.e. the difference between their own AIC and the smallest one, and kept all models with ΔAIC *<* 2. We selected the most parsimonious model from this subset, unless a likelihood ratio test indicated that a more complex one from this subset explained the data significantly better. When discriminating between models with different distributions for the time to the event, we considered the exponential distribution to be the most parsimonious, as it assumed a constant hazard over time.

For both the French and USA datasets, we computed the estimate and standard error of median residence time and colonization rate of each phylogroup according to the AFT models. We defined the median colonization rate as the inverse of the estimated median time to colonization. To obtain estimates of typical colonization rates and residence times for each phylogroup, we computed these estimates for an average host based on those included in the dataset, and for the average values of all the quantitative explanatory factors included in the model. These estimates were used to compute the Spearman’s correlation coefficient between colonization rate and residence time. We used the associated standard errors of parameter estimates to generate confidence intervals by parametric bootstrapping. We randomly sampled these distributions to generate one million sets, including each one residence time and one colonization rate per phylogroup. We computed the Spearman’s correlation coefficient for each set, thus generating a distribution of the correlation between residence and colonization. We considered a correlation to be significant if 95% of the distribution of Spearman’s coefficients was different from zero.

### Correlation between residence and pathogenicity or the number of virulence-associated genes

We used an additional cross-sectional dataset, previously analyzed by Burgaya et al. [49], to estimate pathogenicity and the number of virulence-associated genes (VAG) of each phylogroup. It included 361 commensal and 746 pathogenic clones of *E. coli* (causing bloodstream infections), collected between 2000 and 2017 in Paris and the North of France. These clones were assigned to the same phylogroups as the ones described in the French dataset. These data also included the pathogenic status of each clone (1 for pathogenic, 0 for commensal), as well as the number of VAG carried by the commensal clones, out of a list of 368 genes. Among those genes, some were specifically associated with one of the following type of virulence factors classically associated with pathogenic *E. coli* [46, 47, 68]: toxins (149 genes), adhesin (115 genes), iron acquisition systems (50 genes) and protectins (10 genes).

We estimated the pathogenicity of each phylogroup based on this additional dataset, using a binomial regression with the pathogenic status as a response variable and the phylogroup as the explanatory factor. We used the odds ratio for the association between infection and phylogroup, as estimated from this statistical model, as a proxy for pathogenicity. We also estimated the number of VAG (in total and of each type) using a binomial regression, with the proportion of VAG carried by the clone among those listed as the response variable and the phylogroup as the explanatory factor. We computed the estimated average number of VAG by multiplying the proportion predicted by the model by the total number of genes listed in the data.

As for residence and colonization (see above), we computed the estimates and standard errors for pathogenicity and the numbers of VAG per phylogroup. We used these distributions to generate confidence intervals by parametric bootstrapping. We sampled the distributions to generate one million set of each response variable (pathogenicity, total number of VAG and number of VAG associated with each type of factor) for each phylogroup. We computed a Spearman’s correlation coefficients between each of those variables and the sets of residence times generated earlier (see above). We considered a correlation to be significant if 95% of the distribution of Spearman’s coefficients was different from zero.

### Description of the model of E. coli dynamics

We developed a continuous time, discrete-state Markov chain model describing the dynamics of *E. coli* clones in a host population. In this model, each host could contain between 0 and *N* clones, which belonged to one of *G* possible phylogroups. Hence, the discrete states of the model were the composition of the hosts in *E. coli* clones, expressed as unordered multisets of the phylogroups of the clones they contain. Hosts could be in one of 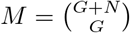 states, which were numbered as follows:

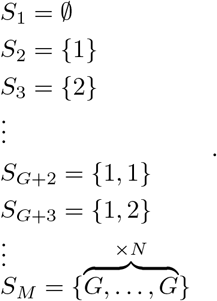

Hence, state *S*_1_ corresponded to a host with no clone, *S*_2_ to one with a clone of phylogroup 1, *S*_*G*+2_ to one with two clones of phylogroup 1, *S*_*M*_ to one with *N* clones of phylogroup *G*. The variables of the model were the set of probabilities *P*_*i*_(*t*) that a host was in state *S*_*i*_ at time *t*. The frequency of phylogroup *k* in the host population at time *t* was defined as:

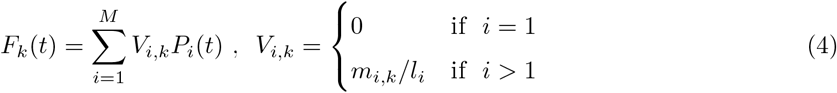

with *m*_*i*,*k*_ the number of clones of phylogroup *k* in state *i* and *l*_*i*_ the total number of clones in state *i. V*_*i*,*k*_ then corresponded to the proportion of clones of phylogroup *k* shed by a host in state *i*.

The transition between states was defined by the matrix **T**_**t**_ of size *M × M*, whose elements, noted *τ*_*i*,*j*_(*t*) *>* 0 for the transition between state *i* and state *j*, were either a clearance rate, noted 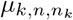, or a colonization rate, noted *λ*_*k*,*n*_(*t*). Both were defined following the formula for an AFT model with an exponential distribution, with a parameter dependence mirroring our findings from the two datasets:

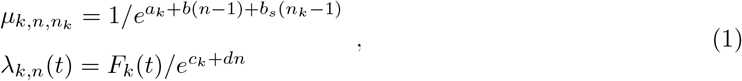

with *a*_*k*_ and *c*_*k*_ phylogroup-specific parameters, *b* and *d* parameters describing the impact of the number of clones in the host *n*, and *b*_*s*_ describing the additional impact of the number of clones of the same phylogroup *n*_*k*_. Competition between clones regardless of phylogroup was expressed either as *b <* 0 if the clearance rate increased with *n* or as *d >* 0 if the colonization rate decreased with *n*. Additional competition between clones of the same phylogroup is expressed as *b*_*s*_ *<* 0, only affecting the clearance rate. The parameters determining these rates were chosen in accordance with the results of the survival analyses.

The transition matrix **T**_**t**_ was sparse, as the transition rate *τ*_*i*,*j*_(*t*) *>* 0 only if state *j* could be reached from state *i* with a single colonization or clearance event. The rows of **T**_**t**_ summed to 0, as the diagonal element 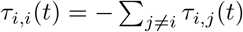. We computed the matrix exponential 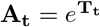 using the package *expm* [69] to get the transition probabilities between states accounting for potential multiple transitions over a single time-step.

We simulated variations in state probabilities in discrete time with a time-step of a day, by updating the elements of the transition matrix **T**_**t**_ using the phylogroup frequencies of the previous day *F*_*k*_(*t−*1). In contrast to clearance rates, colonization rates were indeed not constant over the course of the simulation because they depended on the frequency of each phylogroup in the population. We stopped the simulation when a steady state was reached, i.e. for *t*\* such that 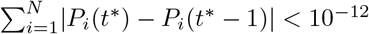. We considered this steady state as the asymptotic state of our system, with the asymptotic probability of state *S*_*i*_ noted 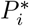.

We used this model to investigate the possible coexistence of several bacterial phylogroups in the population, and the impact of public health measures altering the colonization and clearance rates on the hierarchy of phylogroups. See Supporting information for additional analytical results developped to help interpret the simulation results.

## Acknowledgements

This project has received funding from the CNRS (Momentum grant to F.B.), the French Foundation for Medical Research (Fondation pour la Recherche Médicale, to E.D. and O.Cl) and the European Research Council (ERC) under the European Union’s Horizon 2020 research and innovation program (Grant agreement No. 949208 to F.B.). We thank Loubna Alavoine, Naima Beldjoudi, Charles Burdet, Corinne Da Costa Ribeiro Coutinho, Isabelle Gorenne, Milica Mandic, Ania Mlazindrou, and Valérie Vignali for their help setting up the French clinical study. We thank Cloé Sénéchal for her help with phylogroup typing, and Françoise Chau for her help processing the stool samples. We thank all volunteers for their participation in the studies.

## Author contributions

T.M-J and F.B. designed and conceptualized the project, with inputs from S.L.. R.A., O.Cl., S.D., C.F., J.M., P.R., X.D., F.L.N., S.W., E.D. and F.B. generated the data. T.M-J. analyzed the data with the help of O. Co. and F.B.. T.M-J. and F.B. developed the model, with the help of S.L.. T.M.-J. led the writing of the article, with inputs from all the co-authors.

