## Supporting information for "Residence-colonization trade-off and niche differentiation enable the coexistence of *Escherichia coli* phylogroups in healthy humans"

1. Details of the French dataset
2. Accelerated failure time models
3. Cox models
4. Residence-colonization trade-off among infants
5. Robustness of the results to the number of colonies sampled
6. Analysis of the mathematical model
7. Impact of an external source of *E. coli* clones
8. Supplementary figures
9. Supplementary tables

### 1. Details of the French dataset

We generated the French dataset presented in the main text of the article as part of the EVE/EvoComBac project. This study has been ethically approved by the institutional review board (Comité de Protection des Personnes Ouest IV - Nantes, board approval number 18/20_3). It was sponsored by Assistance Publique - Hôpitaux de Paris. Consents were obtained in compliance with the French law, with signed informed consent obtained from all participants.

From March 2021 to June 2022, we recruited 50 healthy adults in the Paris area to provide stool samples every two weeks (± 3 days) for a planned total of 10 samples. The inclusion criteria were the following:

– healthy volunteer of both sexes between 18 and 65 years inclusive
– subject considered healthy after a thorough general examination (physical exam and questionnaire)
– subject with normal transit with usually one stool per day
– body mass index between 18.5 and 35 kg/m$^2$ inclusive

The non-inclusion criteria were the following:

– subject living in a healthcare institution
– known immunosuppression (HIV, concomitant immunosuppressive treatment, chemotherapy, long-term corticosteroids more than 2 weeks in the last 6 months)
– chronic gastrointestinal diseases (Chronic Ulcerative Colitis, Crohn's Disease)
– any digestive resection except appendectomy and resection of polyps
– subject with a history of bacteremia

Four of the initially fifty enrolled volunteers dropped out of the study without providing any sample and were discarded. Out of the remaining 46 volunteers, 27 were female, 19 were male and the median age at inclusion was of 23 years old. Some volunteers missed a small number of visits, so that we had stool sample for a total of 430 visits. This small level of attrition, in an ecological study of healthy volunteers without intervention, is unlikely to have biased our results. For each visit, we isolated *E. coli* colonies on Drigalski agar plates. Five randomly chosen colonies were sampled for characterization of within-host diversity. This number was chosen as a compromise between quantifying the diversity of dominant types and limiting the total number of samples to be analyzed. Importantly, we chose the colonies to be sampled independently from their phenotype. We used the Clermont method (Clermont et al., 2000, 2013) to assign each colony to a phylogroup. We identified the bacterial species with MALDI-TOF (mass spectrometry) when the results obtained by Clermont typing were incompatible with the species *E. coli*. To differentiate clones within a same phylogroup, we typed all *E. coli* colonies with multiple-locus variable-number tandem-repeat analysis (MLVA, Caméléna et al., 2019). MLVA gave an electrophoretic pattern with 4 to 8 bands per isolate. Two isolates differing by at least one fragment were considered to be from two distinct clones. This level of discrimination was similar to that of the GTG5 Rep-PCR used for the USA dataset (Mohapatra and Mazumder, 2008). Total *E. coli* density was estimated through serial dilutions. Presence of *E. coli* clones over time in each of the 46 hosts in the French dataset is presented in Fig S9.

#### 2. Accelerated failure time models

Accelerated failure time (AFT) models describe how covariates affect the time until the occurrence of an event, here either the colonization or the clearance of a clone. How covariates accelerate or decelerate the occurrence of an event depends on the acceleration $\gamma = exp(-\sum_i \alpha_i X_i)$, with $X_i$ the value of the $i^{th}$ explanatory factor considered and $\alpha_i$ the value of the parameter associated with this factor. The acceleration $\gamma$ correlates positively with the colonization or clearance rate. However, a positive value of $\alpha_i$ indicates a negative impact of the corresponding factor on the rates and a positive impact on the time until the event.

AFT models are parametric, i.e. the time until the event is expected to follow a given distribution. We considered three possible distributions : exponential, Weibull or log-logistic. The exponential distribution is the most parsimonious, as it assumes a constant hazard. Indeed, the other two distributions assume a varying hazard over time, and are characterized by two parameters: the scale, i.e. the inverse of the acceleration factor $\gamma$, which is affected by the covariates, and the shape, which is estimated separately.

##### 2.1. *Selection of AFT models of residence*

We considered the potential impact of the following factors on the residence time, i.e. the time until clearance of each clone:

- the host ($hst$)
- the phylogroup ($grp$)
- the total number of other clones in the host ($ncl$)
- the number of other clones in the host of the same phylogroup as the clone considered ($ngrp$)
- the $\log_{10}$ of the total density of cells in the host ($log(dns)$)

For the analysis of the USA dataset, we also considered the $\log_{10}$ of the density of cells of the clone considered ($log(dscl)$), which was precisely known thanks to the large number of colonies typed.

We fitted AFT models with every possible combination of the model for the distribution of time until clearance, and of these explanatory factors. We computed the AIC of each model and order them by increasing ΔAIC, i.e. the difference between the smallest AIC value and that of the model (see Table S1 and S3 for all models with ΔAIC < 10 for the French and USA data, respectively). We selected the most parsimonious model among those with ΔAIC < 2, unless a likelihood ratio tests showed that another model in this subset explained the data better.

For the French data, two nested models had a ΔAIC < 2:

- **model 1:** $Y \sim hst + grp + ncl + ngrp$
- model 2: $Y \sim hst + grp + ncl + ngrp + log(dns)$

Both models used a log-logistic distribution. We selected model 1 (in bold) because model 2 did not explain the data significantly better (LRT: $\chi^2_{df=1} = 0.650$, $p = 0.4201$).

For the USA data, four models had a ΔAIC < 2. Three of them used an exponential distribution and were nested within one another:

- model 1: $Y \sim hst + grp + ncl + ngrp$
- **model 2:** $Y \sim hst + grp + ncl + ngrp + log(dscl)$
- model 3: $Y \sim hst + grp + ncl + ngrp + log(dns) + log(dscl)$

The last model used a Weibull distribution:

- model 4: $Y \sim hst + grp + ncl + ngrp + log(dscl)$

We selected model 2 (in bold) over the two others using an exponential distribution because it explained data significantly better than model 1 (LRT: $\chi^2_{df=1} = 3.891$, $p = 0.0486$) and as well as model 3 (LRT:

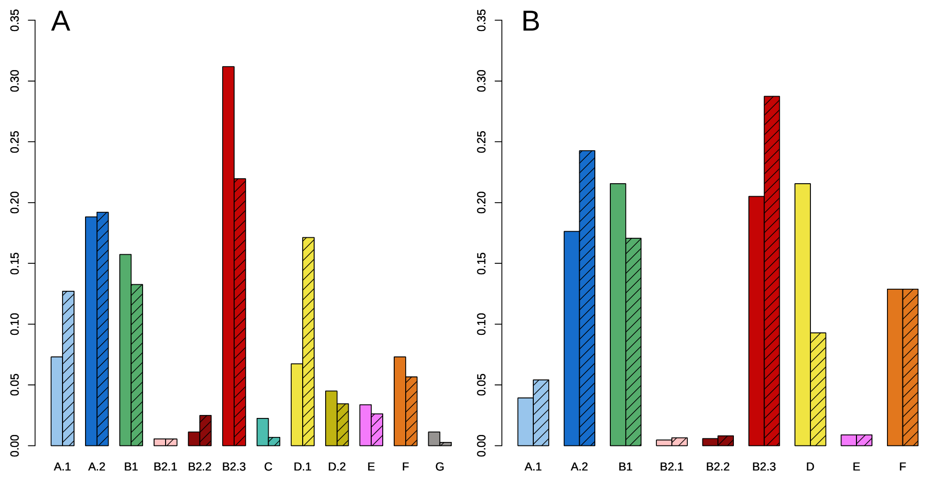


Figure S1: Frequencies of the phylogroups according to the external dataset (clear) and in the dataset itself (hatched), used to correct the colonization rates in the French (A) and in the USA data (B).

$\chi^2_{df=1} = 1.013$, $p = 0.3143$). We also selected model 2 over model 4 because, although they included the same explanatory factors, the former was more parsimonious as it assumed a constant clearance hazard.

Thus, the models best explaining the residence time of *E. coli* in France and USA were very similar. They included the same explanatory factors, except for the addition of the density of the clone considered in the model for the USA dataset, a factor that was not available for the French data.

#### 2.2. *Correction of the colonization rate by the phylogroup frequency*

In addition to the explanatory factors taken into account in the AFT models (see below), we also expected colonization to be impacted by the relative frequencies of the phylogroups in the population. Indeed, more prevalent phylogroups were likely shed in greater quantities, and therefore contributed more to transmission. We assumed as a null model that that phylogroups were shed in proportion to their frequency in the population. Accordingly, we corrected the observed colonization rates by the frequency of the phylogroup in the host population. This was equivalent to multiplying observed colonization times by phylogroup frequency.

We assessed phylogroup frequencies in the host population based on two types of data relevant to the country considered: external data and the dataset itself (Fig S1). For the French dataset, we used cross-sectional external data on the carriage of *E. coli* in healthy hosts of the Paris area, previously analyzed by Burgaya et al. (2023). This dataset is presented in the main text of the article, as it was also used to assess the pathogenicity of the phylogroups and their number of virulence associated genes (see the Material and Methods section). The phylogroups of these 359 clones were defined in the same way as those considered for the French dataset. For the USA dataset, we considered six studies of commensal *E. coli* carriage conducted in the USA (Zhang et al., 2002, Sannes et al., 2004, Johnson et al., 2005, 2007, Hannah et al., 2009, Logue et al., 2012). While they all reported the frequencies of the main phylogroups of *E. coli* (A, B1, B2 and D), they did not necessarily included the minor phylogroups (E and F), which represented 5.1% of the clones in the USA dataset. We therefore assumed that these frequencies corresponded to the ones observed in the USA dataset itself. Besides, these external data did not make the distinction between phylogroups A.1 and A.2 (all noted ‘A’) or between B2.1, B2.2 and B2.3 (all noted ‘B2’). We inferred the relative frequencies of these subgroups based on the USA dataset. For instance, as A.1 and A.2 represented respectively 42% and 58% of the A identified in our dataset, we assigned respectively 42% and 58% of the clones of phylogroup A observed in the external data to A.1 and A.2.

Although the frequencies observed in the external data were comparable to the ones observed in both our datasets (Fig S1), some phylogroups differed in their frequency between the two. To assess whether these differences impacted the negative correlation between residence time and colonization rate observed (see the Results section of the main text of the article), we performed the same analyses using a correction based on the phylogroup frequencies in the dataset themselves. The results were very similar the one presented in the main text, confirming the robustness of this trade-off for the French (95%CI $= [-0.618, -0.832]$, $p = 0.0002$) and USA datasets (95%CI $= [-0.900, -0.583]$, $p < 0.0001$).

#### 2.3. *Selection of AFT models of colonization*

We used the same model selection method as for the AFT models of residence. We considered the potential impact of the following factors on the time until colonization by each clone:

- the host ($hst$)
- the phylogroup ($grp$)
- the total number of other clones in the host ($ncl$)
- the number of other clones in the host of the same phylogroup as the clone considered ($ngrp$)
- the $\log_{10}$ of the total density of cells in the host ($log(dns)$)

We fitted AFT models with every possible combination of the model for the distribution of time until colonization and of these explanatory factors. We computed the AIC of each model and order them by increasing ΔAIC, i.e. the difference between the smallest AIC value and that of the model (see Table S6 and 8 for all models with $\Delta\text{AIC} < 10$ for the French and USA data, respectively). We selected the most parsimonious model among those with $\Delta\text{AIC} < 2$, unless a likelihood ratio tests showed that another model in this subset explained the data better.

For the French data, three nested models had a $\Delta\text{AIC} < 2$:

- **model 1:** $Y \sim hst + grp + ncl$
- model 2: $Y \sim hst + grp + ncl + ngrp$
- model 3: $Y \sim hst + grp + ncl + log(dns))$

All of these models used the Weibull distribution. We selected model 1 (in bold) because neither model 2 (LRT: $\chi^2_{df=1} = 0.721$, $p = 0.3957$) nor model 3 (LRT: $\chi^2_{df=1} = 0.456$, $p = 0.4994$) explained the data significantly better.

For the USA data, four nested models had a $\Delta\text{AIC} < 2$:

- **model 1:** $Y \sim hst + grp + ncl$
- model 2: $Y \sim hst + grp + ncl + ngrp$
- model 3: $Y \sim hst + grp + ncl + log(dns)$
- model 4: $Y \sim hst + grp + ncl + ngrp + log(dns)$

All of these models used the Weibull distribution. We selected model 1 (in bold) because neither model 2 (LRT: $\chi^2_{df=1} = 1.815$, $p = 0.1779$), model 3 (LRT: $\chi^2_{df=1} = 0.650$, $p = 0.4203$) nor model 4 (LRT: $\chi^2_{df=2} = 2.534$, $p = 0.2817$) explained the data significantly better. As for the models of residence, the models best explaining the colonization rates in France and the USA were consistent. Indeed, the explanatory factors included were the same for both datasets.

#### 3. Cox models

To confirm the results of the survival analyses using the AFT models (see the Results section of the main text of the article), we compared them to those of semi-parametric Cox models including the same explanatory factors. These models compute hazard ratios indicating the relative risk of events – colonization or clearance – occurring depending depending on the factors considered. Unlike AFT models, Cox models make no assumptions about the shape of the hazard over time, but assume proportional hazards between strata. In those models, a factor increasing the hazard decreases the time until the event. Therefore, positive coefficients in one type of model corresponds to negative coefficients in the other.

The host factor systematically violated the proportional hazard hypothesis, for residence (French dataset: $\chi^2_{fd=45} = 123.3161$, $p < 0.0001$; USA dataset: $\chi^2_{fd=7} = 24.75$, $p = 0.0008$) and for colonization (French dataset: $\chi^2_{fd=7} = 139.1213$, $p < 0.0001$; USA dataset: $\chi^2_{fd=7} = 51.70$, $p < 0.0001$). Therefore, we used Cox models stratified by hosts. Otherwise, we included the same explanatory factors as those used in the corresponding best AFT models.

The Cox models provided results consistent with the AFT models (Table S11-S14). The sign and significance of the coefficients associated with quantitative factors were consistent. Although the significance of the coefficients associated to the phylogroups was sometimes different in the Cox model, their sign was also always consistent with that of the AFT models. These complementary analyses therefore confirm our initial observations.

#### 4. Residence-colonization trade-off among infants

We assessed whether the trade-off between residence and colonization identified among adults (see the Results section of the main text of the article) was also observed during the initial colonization of the gut of infants. To do so, we used an additional longitudinal dataset, previously analyzed by Ostblom et al. (2011). It included longitudinal samples in 130 infants at fixed times since birth (after 3 days, 1, 2 and 4 weeks, 2, 6 and 12 months). The 273 clones of *E. coli* identified in these samples were assigned to one of the four main *E. coli* phylogroups (A, B1, B2 and D).

We used accelerated failure time (AFT) models to analyze these data, using the same method, explanatory factors and time distributions as described above (see Supporting information 2). To correct the observed colonization rates, we used the phylogroup frequencies in the dataset itself. No external study provided the frequency of *E. coli* phylogroups in healthy infants in Sweden, as the other published studies were based on the same cohort (Nowrouzian et al., 2005, Karami et al., 2007)).

We used the same model selection method as presented in Supporting information 2, based on AIC (see Table S5 and S10 for all models with $\Delta$AIC $< 10$ for residence and colonization, respectively). Four nested AFT models explaining the residence time had a $\Delta$AIC $< 2$:

- **model 1:** $Y \sim grp + ncl$
- model 2: $Y \sim grp + ncl + log(dns)$
- model 3: $Y \sim grp + ncl + log(dncl)$

All these models used a Weibull distribution. We selected model 1 (highlighted in bold) because neither model 2 (LRT: $\chi^2_{df=1} = 0.065$, $p = 0.7984$), model 3 (LRT: $\chi^2_{df=1} = 0.190$, $p = 0.6628$) nor model 4 (LRT: $\chi^2_{df=1} = 0.135$, $p = 0.7135$) explained the data significantly better. The number of clones in the same phylogroup was not a factor in the model that best explained residence time for this data set, in contrast to the best AFT models for French and American data.

Three nested AFT models explaining the colonization rate had a $\Delta$AIC $< 2$:

- **model 1:** $Y \sim grp + ncl$
- model 2: $Y \sim grp + ncl + ngrp$

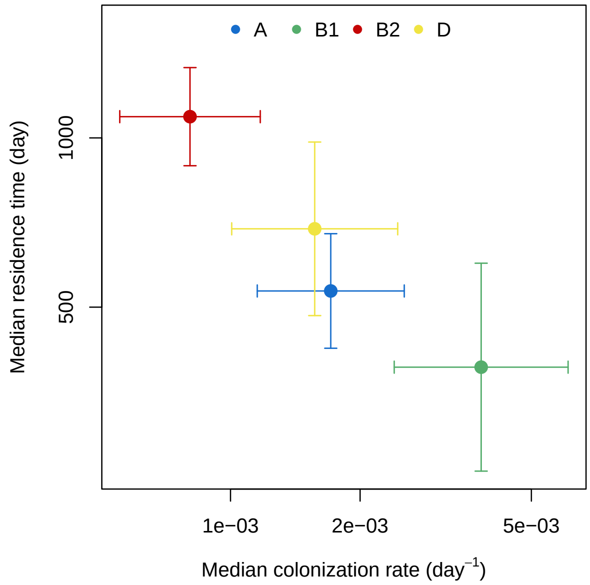


Figure S2: Median residence time as a function of the median colonization rate, as estimated for each phylogroup in the infant dataset, for an average number of clones. Dots represent estimates and bars their 95% confidence intervals.

– model 3: $Y \sim grp + ncl + log(dns)$

These models all used the Weibull distribution. Again, we selected model 1 because neither model 2 (LRT: $\chi^2_{df=1} = 0.085$, $p = 0.7707$) nor model 3 (LRT: $\chi^2_{df=1} = 0.263$, $p = 0.6079$) explained the data significantly better. The host factor was included in neither models best explaining residence or colonization. This was likely due to the very large number of infants (130) which reduced considerably the parsimony of models including this explanatory factor.

The selected models provided estimates and standard errors for the residence times and colonization rates (Fig S2). We observed a trade-off between the colonization and residence abilities of *E. coli* phylogroups during the initial colonization of the gut of infants during their first year. The negative correlation was significant (parametric bootstrapping; Spearman's $\rho = -0.969$, 95%CI = $[-0.996, -0.496]$, $p = 0.0011$).

### 5. Robustness of the results to the number of colonies sampled

While the French dataset encompassed a larger number of subjects, the number of colonies used for each sample (five) was lower than that of the USA dataset, which was of the order of 95 (Martinson et al., 2019). We tested how this difference affected our ability to detect the intra-host diversity of clones within the samples, and the conclusions of our AFT models.

We compared the diversity observed in the two datasets. Diversity was computed as the number of distinct clones per sample. The diversity was comparable, and slightly higher in the French population, in spite of less exhaustive sampling. Diversity was 1.74 clones per sample in France, compared with 1.56 in the USA (Fig S3A). Geographical differences in clone diversity could explain the slightly higher diversity in France (Escobar-Páramo et al., 2004, Skurnik et al., 2008). Yet, the distribution of the number of clones dropped rapidly in both datasets, indicating that colonization by more than five clones was rare.

We assessed how diversity could be reduced by the typing of a limited number of colonies. We sub-sampled the USA dataset by randomly selecting five colonies by sample without replacement. Therefore, the probability that a clone appeared in a sub-sampled dataset was proportional to the fraction of colonies of this clone in the sample. Sub-sampling reduced the intra-host diversity to 1.24 clone per sample (Fig S3B). This confirms that the diversity of *E. coli* in France is even larger than suggested by Fig S3A.

We then investigated how this sub-sampling could alter our results on niche differentiation and the colonization-residence trade-off. We performed survival analyses for residence and colonization on the sub-sampled datasets, using the same time distribution and explanatory factors as for the analysis of the complete data. The impact of the number of clones of the same phylogroup was only significant, i.e. with a p-value $< 0.05$, in 10.3% of the AFT models, presumably due to the more limited diversity caused by sub-sampling. However, there was no evidence for a bias in the estimates of this parameter when diversity was reduced. Indeed, the model estimate for the complete data (coef $= -1.108$) was in the $44^{th}$ percentile of the distribution of estimates for the sub-sampled data (Fig S4).

We also computed the predicted median residence time and colonization rate of each phylogroup from the AFT models on all the subsets, and measured their correlation. Despite the sub-sampling, the estimates were overall similar to those based on the complete USA dataset (Fig S5). Main differences

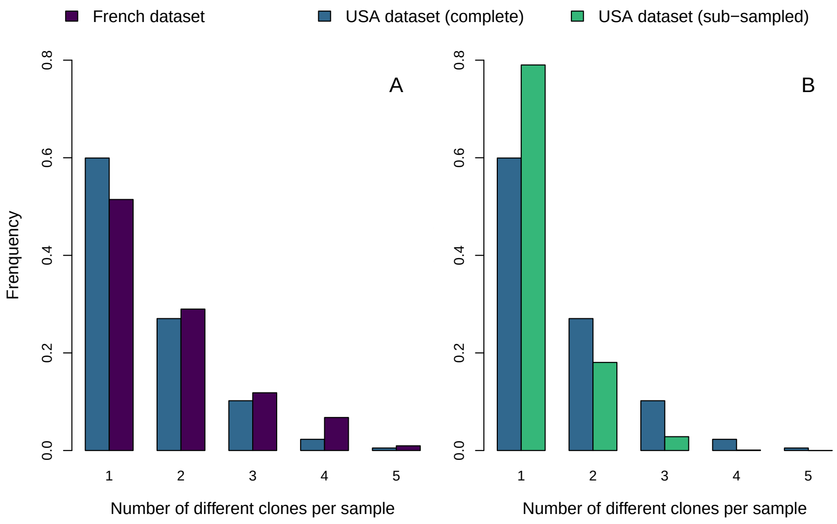


Figure S3: Distribution of the number of different clones per sample in the French dataset (purple), USA dataset (blue) and the subsets of the USA dataset created by randomly selecting five colonies by sample (green). A compares the diversity between the two datasets. B compares the diversity in the USA dataset with and without sub-sampling.

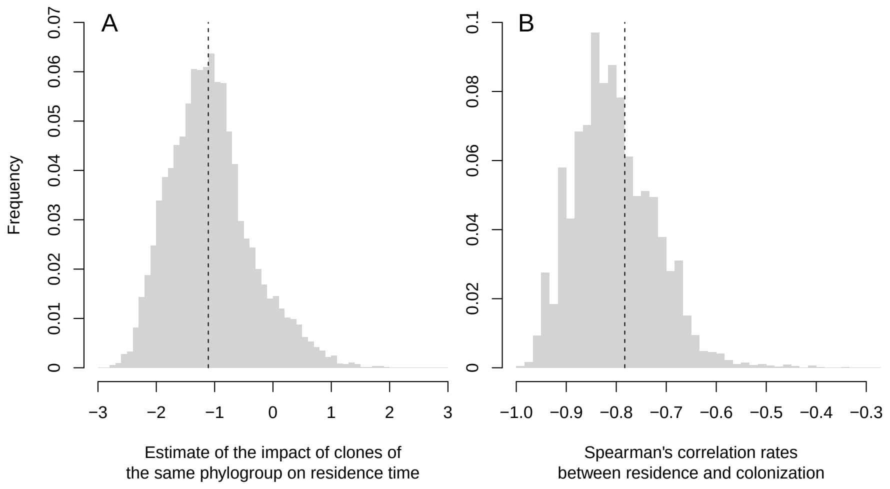


Figure S4: Distribution of the estimates of the impact of the number of clones of the same phylogroups on the residence time of the clones (A) and of the Spearman's correlation between residence time and colonization rate (B), for the sub-sampled USA dataset (grey) and for the complete dataset (dashed line).

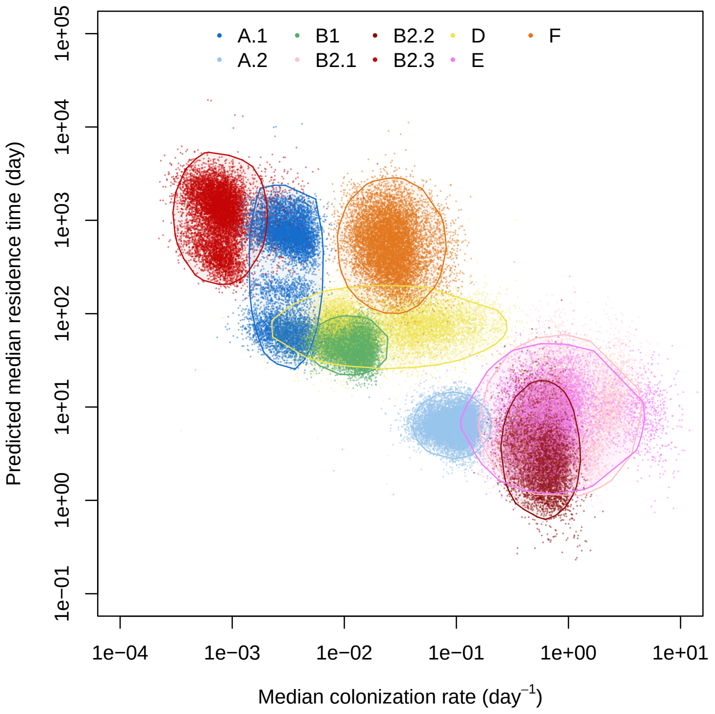


Figure S5: Median residence time as a function of the median colonization rate for each phylogroup according to analyses of the reconstructed datasets. Dots represent individual estimates from the analysis of each dataset. Polygons encompass 95% of the estimates for a given phylogroup.

concerned the phylogroup A and D, whose estimated survival were respectively lower and higher. However, the correlation between colonization and residence remained largely negative for the reconstructed datasets, with 95% of the distribution of Spearman's coefficients between $-0.93$ and $-0.63$. Once again, there was no strong bias induced by sub-sampling, as the correlation rate for the complete data (Spearman's $\rho = -0.733$) was in the $64^{th}$ percentile of the distribution.

### 6. Analysis of the mathematical model

This section presents the analysis of the mathematical model describing *E. coli* colonization dynamics presented in the main text. Firstly, we present the model, which describes how the colonization status of hosts changes by the events of colonization and clearance. Secondly, we examine what happens when a single bacterial phylogroup is present in the host population. We show how the invasion dynamics of the phylogroup in an 'empty' (non-colonized) host population depends on the colonization-to-clearance ratio $R_k$, which is also the basic reproduction number in epidemiology. We derive the equilibrium endemic distribution of the number of clones of this phylogroup, and the dependence on competition parameters. Thirdly, we consider a case with two phylogroups. We show that under some conditions on competition (symmetric competition among phylogroups, no niche differentiation), a new phylogroup can invade the population at endemic equilibrium if and only if its colonization-to-clearance ratio $R_k$ is greater than that of the resident strain. Finally, we show how changes in transmission or clearance can alter the competitive hierarchy between phylogroups.

#### 6.1. *Presentation of the model*

In our model, a clone of phylogroup $k$ in a host with $n$ colonizing strains is characterized by a clearance rate $\mu_{k,n,n_k}$, such that:

$$\mu_{k,n,n_k} = 1/e^{a_k+b(n-1)+b_s(n_k-1)} = e^{-a_k}\ e^{-b(n-1)}e^{-b_S(n_k-1)}. \tag{1}$$

When there is no niche differentiation creating additional clearance of clones of the same phylogroup, i.e. $b_S = 0$, the clearance rate can be expressed as:

$$\mu_{k,n,n_k} = e^{-a_k}\ e^{-b(n-1)}. \tag{2}$$

We expect the coefficient $b$ to be negative if more clones lead to a higher clearance rate, with $e^{-a_k}$ corresponding to the clearance rate in hosts with a single clone.

The clone is also characterized by a colonization rate $\lambda_{k,n}(t)$, such that:

$$\lambda_{k,n}(t) = F_k(t)/e^{c_k+dn} = F_k(t)e^{-c_k}e^{-dn}. \tag{3}$$

We expect coefficient $d$ to be positive if more clones lead to lower colonization rates. In contrast to the clearance rate, the colonization rate of phylogroup $k$ depends on its frequency $F_k(t)$ in the host population, defined as:

$$F_k(t) = \sum_{j=1}^{M} V_{j,k}\ P_j(t), \tag{4}$$

with $P_j(t)$ the frequency of state $j$, and $V_{j,k}$ the frequency of phylogroup $k$ in host state $j$. We consider here that the phylogroup is transmitted proportionally to its frequency in each host state, i.e. $V_{j,k} = m_{j,k}/l_j$ with $m_{j,k}$ the number of clones of phylogroup $k$ and $l_j$ the total number of clones in state $j$. As a convention, $V_{1,k} = 0$, as state 1 corresponds to an empty host.

#### 6.2. *Model with a single phylogroup*

Let us consider a single phylogroup $k$ ($G = 1$), such that the frequency of hosts colonized by $n$ clones of this phylogroup is noted $X_{k,n}(t)$. All frequencies sum to 1, i.e. $\sum_{n=0}^{\infty} X_{k,n}(t) = 1$. The dynamics of the frequencies of hosts can be expressed as:

$$\begin{aligned}
\dot{X}_{k,0} &= -\lambda_{k,0}(t)X_{k,0} + \mu_{k,1,1}X_{k,1} \\
\dot{X}_{k,n} &= \lambda_{k,n-1}(t)X_{k,n-1} - \lambda_{k,n}(t)X_{k,n} + \mu_{k,n+1,n+1}X_{k,n+1} - \mu_{k,n,n}X_{k,n} \quad \forall n \geq 1
\end{aligned} \tag{5}$$

In the first equation, the first term represents colonization of empty hosts, and the second term clearance from hosts with one clone. In the second equation, the first and second terms respectively represent colonization of hosts with $n-1$ and $n$ clones, the third and fourth terms respectively represent clearance of hosts with $n-1$ and $n$ clones. The time dependency of the variables $X_{k,n}$ is omitted for clarity. Replacing the expressions for $\mu_{k,n,n}$ and $\lambda_{k,n}(t)$ according Eq 2 and Eq 3 respectively give:

$$\begin{aligned}
\dot{X}_{k,0} = &\quad -F_k(t)e^{-c_k}e^{-d\times 0}X_{k,0} + e^{-a_k}e^{-b\times 0}X_{k,1} \\
\dot{X}_{k,n} = &\quad F_k(t)e^{-c_k}e^{-d(n-1)}X_{k,n-1} - F_k(t)e^{-c_k}e^{-dn}X_{k,n} + e^{-a_k}e^{-bn}X_{k,n+1} \\
&\quad -e^{-a_k}e^{-b(n-1)}X_{k,n} \quad \forall n \geq 1
\end{aligned} \tag{6}$$

We can rescale time by $e^{-a_k}$ without loss of generality, such that the mean residence time of the phylogroup $k$ in a host colonized by a single clone is 1. The dynamics described in Eq 6 then becomes:

$$\begin{aligned}
\dot{X}_{k,0} &= -F_k(t)\tfrac{e^{-c_k}}{e^{-a_k}}e^{-d\times 0}X_{k,0} + e^{-b\times 0}X_{k,1} \\
\dot{X}_{k,n} &= F_k(t)\tfrac{e^{-c_k}}{e^{-a_k}}e^{-d(n-1)}X_{k,n-1} - F_k(t)\tfrac{e^{-c_k}}{e^{-a_k}}e^{-dn}X_{k,n} + e^{-bn}X_{k,n+1} - e^{-b(n-1)}X_{k,n} \quad \forall n \geq 1
\end{aligned} \tag{7}$$

Defining the colonization-to-clearance ratio, i.e. the basic reproduction number of phylogroup $k$, $R_k := e^{a_k - c_k}$, Eq 7 gives:

$$\begin{aligned}
\dot{X}_{k,0} &= -F_k(t)R_k e^{-d\times 0}X_{k,0} + e^{-b\times 0}X_{k,1} \\
\dot{X}_{k,n} &= F_k(t)R_k e^{-d(n-1)}X_{k,n-1} - F_k(t)R_k e^{-dn}X_{k,n} + e^{-bn}X_{k,n+1} - e^{-b(n-1)}X_{k,n} \quad \forall n \geq 1
\end{aligned} \tag{8}$$

##### 6.2.1 Invasion dynamics of the phylogroup when rare depends on the colonization-to-clearance ratio $R_k$

Here we show that the dynamics of a newly introduced phylogroup in the host population depends on the colonization-to-clearance ratio $R_k$. Let us consider a case where phylogroup $k$ is the only one present in the host population at low prevalence, i.e. $X_{k,0}$ is close to 1, $X_{k,1}$ is close to 0 and $X_{k,n} = 0$ for $n > 1$. As above, the time-dependency of the variables $X_{k,n}$ is omitted for clarity. In this case, $F_k$ is also close to 0. More precisely, $F_k(t) = X_{k,1} = 1 - X_{k,0}$ under our assumptions for transmission. The system then reduces to the first order to:

$$\begin{aligned}
\dot{X}_{k,0} &= -F_k(t)R_k e^{-d\times 0}X_{k,0} + e^{-b\times 0}X_{k,1} \\
\dot{X}_{k,1} &= F_k(t)R_k X_{k,0} - X_{k,1} \quad \forall n \geq 1
\end{aligned} \tag{9}$$

Then, phylogroup $k$ increases in frequency if and only if $\dot{X}_{k,1} > 0$, meaning that:

$$F_k(t)R_k X_{k,0} - X_{k,1} = X_{k,1}R_k X_{k,0} - X_{k,1} \approx X_{k,1}\left(R_k - 1\right) > 0. \tag{10}$$

This condition is equivalent to $R_k > 1$. Therefore, the ability of phylogroup $k$ to invade a population is defined by its colonization-to-clearance ratio $R_k$.

##### 6.2.2 Endemic equilibrium solutions

Once a phylogroup is introduced and increases in prevalence, it will reach a non-zero prevalence with an equilibrium distribution of the multiplicity of colonization. Here we find the equilibrium solution for the frequency of each multiplicity of colonization when $R_k > 1$. The equilibrium value for $X_{k,1}$ verifies:

$$\widehat{X}_{k,1} = F_k(t)\ R_k\ \widehat{X}_{k,0}. \tag{11}$$

The next value can be found by recursion, solving for $\dot{X}_{k,n} = 0$:

$$\widehat{X}_{k,n+1} = F_k(t)\ R_k\ \left(e^{(b-d)n}\widehat{X}_{k,n} - e^{b\ n-d\ (n-1)}\widehat{X}_{k,n-1}\right) + e^b\ \widehat{X}_{k,n}. \tag{12}$$

For $n = 1$, Eq 12 gives:

$$\widehat{X}_{k,2} = F_k(t)\ R_k\ \left(e^{(b-d)}\widehat{X}_{k,1} - e^b\widehat{X}_{k,0}\right) + e^b\ \widehat{X}_{k,1}. \tag{13}$$

Replacing $\widehat{X}_{k,1}$ according to Eq 11 gives:

$$\widehat{X}_{k,2} = F_k(t)\ R_k\ \left(e^{(b-d)}F_k(t)\ R_k\ \widehat{X}_{k,0} - e^b\ \widehat{X}_{k,0}\right) + e^b\ F_k(t)\ R_k\ \widehat{X}_{k,0} \tag{14}$$

Using the ansatz $\widehat{X}_{k,n} = a_n\left[F_k(t)\ R_k\right]^n\ \widehat{X}_{k,0}$ (with $a_1 = 1$), replacing in Eq 12 gives:

$$\begin{aligned}\widehat{X}_{k,n+1} = {} & F_k(t)\ R_k\ \left(e^{(b-d)n}\ a_n\ \left[F_k(t)\ R_k\right]^n\ \widehat{X}_{k,0} - e^{b\ n-d\ (n-1)}a_{n-1}\left[F_k(t)\ R_k\right]^{n-1}\ \widehat{X}_{k,0}\right) + \\ & e^b\ a_n\ \left[F_k(t)\ R_k\right]^n\ \widehat{X}_{k,0} \\ \widehat{X}_{k,n+1} = {} & \left(e^{(b-d)n}\ a_n\ \left[F_k(t)\ R_k\right]^{n+1}\ \widehat{X}_{k,0}\right) + \left(e^b\ a_n - e^{b\ n-d\ (n-1)}a_{n-1}\right)\ \left[F_k(t)\ R_k\right]^n\ \widehat{X}_{k,0}\end{aligned}. \tag{15}$$

From the first term, $a_{n+1}$ must satisfy the following recursion:

$$a_{n+1} = e^{(b-d)n}\ a_n. \tag{16}$$

In the second term, replacing $a_n$ by $e^{(b-d)(n-1)}\ a_{n-1}$ gives:

$$\left[e^b\ e^{(b-d)(n-1)} - e^{b\ n-d\ (n-1)}\right]\ a_{n-1}. \tag{17}$$

The coefficient in brackets cancels, so that the solution is of the form:

$$\widehat{X}_{k,n} = a_n\ \left[F_k(t)\ R_k\right]^n\ \widehat{X}_{k,0}. \tag{18}$$

The series is therefore as follows:

$$\begin{aligned} & a_1 = 1 \\ & a_2 = e^{(b-d)} \\ & a_3 = e^{2(b-d)}e^{(b-d)} \\ & a_4 = e^{3(b-d)}e^{2(b-d)}e^{(b-d)} \\ & a_n = e^{(b-d)\sum_{i=1}^{n-1} i} = e^{(b-d)\sum_{i=1}^{n-1} i} = e^{(b-d)n(n-1)/2}\end{aligned}. \tag{19}$$

Therefore, the general solution for $\widehat{X}_{k,n}$ is as follows:

$$\widehat{\mathbf{X}}_{k,n} = e^{(\mathbf{b}-\mathbf{d})\frac{n(n-1)}{2}}\left[F_k(t)R_k\right]^n\widehat{X}_{k,0} \tag{20}$$

The frequency of colonized hosts decreases for large enough values of $n$ if $(b-d) < 0$ and also decreases for any $n$ if $F_k(t)R_k < 1$.

The phylogroup frequency $F_k(t)$ is a linear combination of $\widehat{X}_{k,0}$. Under our assumption for transmission, $F_k(t) = 1 - \widehat{X}_{k,0}$ as host can only carry clones of phylogroup $k$. Eq 20 therefore becomes:

$$\widehat{\mathbf{X}}_{k,n} = e^{(\mathbf{b}-\mathbf{d})\frac{n(n-1)}{2}} {R_k}^n \left(1 - \widehat{X}_{k,0}\right)^n \widehat{X}_{k,0} \tag{21}$$

Solving for $\widehat{X}_{k,0}$ with the constraint that all host frequencies must sum to 1 to obtain an explicit expression, Eq 21 gives:

$$\sum_{n=0}^{\infty} \widehat{X}_{k,n} = 1 = \widehat{X}_{k,0} \sum_{n=0}^{\infty} e^{(b-d)\frac{n(n-1)}{2}} R_k^n \left(1 - \widehat{X}_{k,0}\right)^n \tag{22}$$

A trivial solution is $\widehat{X}_{k,0} = 1$, i.e. every clone is cleared and the phylogroup does not persist. The non-trivial solution in $[0, 1)$ does not have an explicit solution, as the series in $\sum_{n=1}^{\infty} e^{-n^2} A^n$ converges but does not have an explicit solution. For $\widehat{X}_{k,0}$ close to 0, the right-hand side of Eq 22 is approximately linear in $X_{k,0}$ with a positive slope of value $\sum_{n=0}^{\infty} e^{(b-d)\frac{n(n-1)}{2}} R_k^n$. For $\widehat{X}_{k,0}$ close to 1, the series is approximately $1 + (R_k - 1)(1 - \widehat{X}_{k,0})$ and decreases linearly with $\widehat{X}_{k,0}$. If there is only one value of $\widehat{X}_{k,0} \in [0, 1($ for which the series equals 1, then the series is an increasing function of $\widehat{X}_{k,0}$ at this solution. Therefore, $\widehat{X}_{k,0}$ is a decreasing function of $R_k$.

#### 6.3. *Model with multiple phylogroups*

We now investigate what happens when an invading phylogroup is introduced in the population of the resident phylogroup at equilibrium. In the following, we show that $R_k$ is the fitness quantity determining whether the invading phylogroup will rise in frequency or not. Let us consider a 'resident' phylogroup $k$ established at endemic equilibrium and an 'invading' phylogroup $v$ at low density in the host population. The system governing the dynamics of the phylogroups can be described as:

$$\begin{aligned}
\dot{X}_{k,n} = \; & F_k(t)\ R_k\ e^{-d\ (n-1)} X_{k,n-1} - \left( R_k F_k(t) + \underbrace{R_v F_v(t)}_{\text{colonization by } v} \right) e^{-d\ n} X_{k,n} \\
& + e^{-b\ n} \left( X_{k,n+1} + \underbrace{\frac{Y_{v,n+1}}{n+1}}_{\text{clearance of } v} \right) - e^{-b(n-1)}\ X_{k,n} \quad \forall n \geq 1 \\
\dot{Y}_{v,n} = \; & e^{-d\ (n-1)} \left( \underbrace{F_k(t)\ R_k\ Y_{v,n-1}}_{\text{colonization by } k} + \underbrace{F_v(t)\ R_v\ X_{k,n-1}}_{\text{colonization by } v} \right) - \underbrace{F_k(t)\ R_k\ e^{-d\ n} Y_{v,n}}_{\text{colonization by } k} \\
& + \underbrace{e^{-b\ n}\ \frac{n}{n+1}\ Y_{v,n+1}}_{\text{clearance of } k} - e^{-b(n-1)}\ Y_{v,n}\ \forall n \geq 1
\end{aligned} \quad , \tag{23}$$

with $Y_{v,n}$ the density of hosts carrying one clone of phylogroup $v$ and $(n-1)$ clones of phylogroup $k$ $\forall\ n \geq 1$, $R_v$ the reproduction number of phylogroup $v$ (Fig S6) and $F_v(t)$ is the frequency of phylogroup $v$.

The first equation differs from the previous equation describing a monomorphic population by two terms. First, the colonization by the phylogroup $v$ reduces the frequency of hosts colonized with $n$ strains of phylogroup $k$ at a rate proportional to $R_v F_v(t)$. Second, hosts of type $(v, n+1)$ become of type $(k, n)$ when losing their clone of phylogroup $v$. In the second equation, hosts of type $(v, n-1)$ or of type $(k, n-1)$ become of type $(v, n)$ when colonized by a clone of phylogroup $k$ and phylogroup $v$, respectively. Furthermore, hosts of type $(v, n+1)$ also become of type $(v, n)$ with the clearance of a clone of phylogroup $k$. Here, we assume that colonization by multiple clones of phylogroup $v$ is negligible, as the frequency of this phylogroup is much smaller than that of phylogroup $k$. We can deduce the proportion of empty

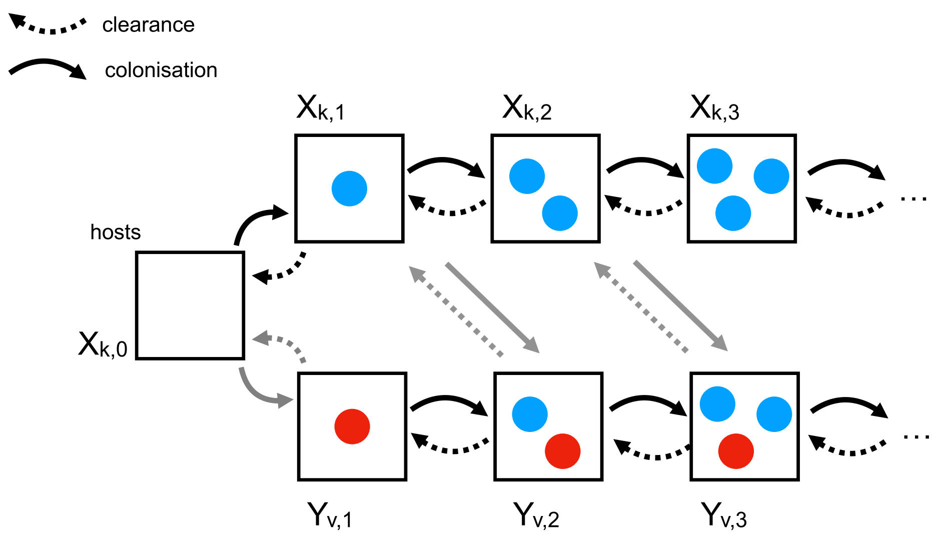


Figure S6: Potential transitions between host states (squares) because of clearance or colonization of either resident clones (blue dot) or invading clones (red dot). States with more than one invading clone are neglected. Arrows correspond to colonization (solid line) or clearance (dashed line) of clones of the resident (black line) or invading (grey line) phylogroup.

hosts from the condition that $\sum_{i=0}^{\infty} X_{k,i} + \sum_{i=1}^{\infty} Y_{v,i} \approx \sum_{i=0}^{\infty} X_{k,i} = 1$.

The equilibrium with phylogroup $k$ alone corresponds to $X_{k,n} = \widehat{X}_{k,n}$ and $Y_{v,n} = 0 \ \forall n \geq 1$. The dynamics of the system linearized at this equilibrium are given by the matrix:

$$\begin{bmatrix} \mathbf{J}_k & \mathbf{S} \\ \mathbf{0} & \mathbf{J}_v \end{bmatrix}.$$

The matrix $\mathbf{J_k}$ contains the partial derivatives of $\dot{X}_{k,n}$ with respect to $X_{k,n}$, evaluated at the monomorphic equilibrium $\{\widehat{\mathbf{X}}_k, \mathbf{0}\}$. $\widehat{\mathbf{X}}_k$ corresponds to the vector of $\widehat{X}_{k,n}$ values $\forall\ n \geq 1$ and $\mathbf{0}$ the vector of 0s, i.e. the equilibrium for $Y_{v,n}$. By definition of the stable equilibrium, the spectral bound of this matrix $\mathbf{J_k}$ is smaller than 0 (Hurford et al., 2010). The matrix $\mathbf{J_v}$ contains the partial derivatives of $\dot{Y}_{v,n}$ with respect to $Y_{v,n}$, evaluated at the monomorphic equilibrium $\{\widehat{\mathbf{X}}_k, \mathbf{0}\}$. The matrix $\mathbf{S}$ contains the partial derivatives of $X_{k,n}$ with respect to $Y_{v,n}$, evaluated at the monomorphic equilibrium $\{\widehat{\mathbf{X}}_k, \mathbf{0}\}$. The lower-left matrix is $\mathbf{0}$ because the derivatives of $\dot{Y}_{v,n}$ with respect to $X_{k,n}$ evaluated at the invader-free equilibrium are all 0. The stability of the equilibrium is therefore governed by the spectral bound of $\mathbf{J_v}$, noted $s(\mathbf{J_v})$.

Phylogroup $v$ is at a neutral equilibrium, $s(\mathbf{J_v}) = 0$ if it has the same characteristics as $k$, i.e. if $R_v = R_k$ (Hurford et al., 2010). The two phylogroups differ only in their basic reproduction number $R_k$ and $R_v$, as the competition coefficient $b$ and $d$ are assumed to be identical for both phylogroups. Thus, phylogroup $v$ can invade the host population if $R_v > R_k$, as $s(\mathbf{J_v}) > 0$ in this case. Conversely, it cannot invade if $R_v < R_k$ as then $s(\mathbf{J_v}) < 0$.

#### 6.4. *Impact of changes in colonization or clearance rates on bacterial evolution*

We now investigate how non-phylogroup-specific changes affect the competitive relationship between phylogroups. Let us consider two phylogroups $x$ and $y$ so that $R_x > R_y$. We first consider a 'multiplicative' change, so that the clearance or colonization rate of every phylogroup is multiplied by a strictly positive constant factor, e.g. by defining new clearance rates $\mu'_{k,n,n_k} = \eta \times \mu_{k,n,n_k}$, leading to $R'_k = R_k/\eta$. Because $\eta$ is strictly positive ($\eta > 1$ and $\eta < 1$ respectively correspond to an increase and a decrease of the clearance rate), $R_x^{'} > R_y^{'}$ for any value of $\eta$. Therefore, the hierarchy between phylogroups remains unaffected by such changes. Modifying colonization rate instead of clearance would lead to the same result, as multiplying the clearance rate by $\eta$ has the same impact on the colonization-to-clearance ratio as multiplying the colonization rate by $1/\eta$.

Then, we consider an 'additive' change, so that a constant factor is added to the clearance or colonization rate of every phylogroup, e.g. by defining new clearance rates $\mu_{k,n,n_k}^{''} = \mu_{k,n,n_k} + \theta$. This change is tantamount to multiplying the clearance rate by a phylogroup-specific factor $\theta_k^m$:

$$\mu_{k,n,n_k}^{''} = \theta_k^m \mu_{k,n,n_k},\ \theta_k^m = \frac{\mu_{k,n,n_k} + \theta}{\mu_{k,n,n_k}}, \tag{24}$$

leading to $R_k^{''} = R_k/\theta_k^m$. Following Eq 24, this factor $\theta_k^m$ is negatively correlated with $\mu_{k,n,n_k}$, meaning that this change disproportionately affects phylogroups with a slow turnover. Therefore, there is a factor $\theta^{eq}$ corresponding to the additional clearance rate canceling out the difference between $R_x$ and $R_y$:

$$\theta^{eq} = \frac{R_x - R_y}{R_y e^{a_x} - R_x e^{a_y}}. \tag{25}$$

The phylogroup hierarchy is maintained, i.e. $R_x^{''} > R_y^{''}$ if $\theta < \theta^{eq}$, but reversed, i.e. $R_x^{''} < R_y^{''}$ if $\theta > \theta^{eq}$. The value of $\theta^{eq}$ is positive only if $a_x < a_y$, meaning that an additional source of clearance ($\theta > 0$) can only change the phylogroup hierarchy if the phylogroup with the higher initial colonization-to-extinction rate $R_k$ also has the lower initial clearance rate $\mu_{k,n,n_k}$.

### 7. Impact of an external source of *E. coli* clones

Our model assumes that any colonizing clone is directly transmitted from another host in the population. This implies two hypotheses: that transmission of *E. coli* is solely human-to-human, and that the host population is completely isolated. The first hypothesis ignores the potential role of the environment, which harbors various strains of *E. coli* (Money et al., 2010, Ludden et al., 2019) also susceptible to be transmitted to humans. The second hypothesis would be invalidated by rare transmissions of *E. coli* from hosts belonging to another population. Both hypotheses can be relaxed by modeling colonization as depending on a mixture of human-to-human within-population transmission and transmission from an external source also providing clones.

Following Levin et al. (Levin et al., 1997), we considered a new colonization rate $\lambda^r_{k,n}(t)$, which depended in proportion $(1-q)$ on transmission from other hosts in the population, and in proportion $q$ on influx from this constant reservoir:

$$\lambda^r_{k,n}(t) = \frac{qF^r_k + (1-q)\,F_k(t)}{e^{c_k+dn}},$$

with $F^r_k$ the frequency of phylogroup $k$ in the reservoir. This rate can also be expressed as the sum of two rates respectively corresponding to transmission from the reservoir and transmission within the population:

$$\lambda^r_{k,n}(t) = \overbrace{qF^r_k e^{-(c_k+dn)}}^{\text{reservoir}} + \overbrace{(1-q)\,F_k(t)e^{-(c_k+dn)}}^{\text{within population}},$$

with the within-population part of the colonization rate corresponding to $(1-q)\,\lambda_{k,n}(t)$. By definition, the contribution of the reservoir corresponds to a minimum colonization rate for phylogroup $k$, i.e. $\lambda^r_{k,n}(t) \geq qF^r_k e^{-(c_k+dn)}$. This threshold is always positive for any value of $q > 0$ and $F^r_k > 0$. Hence, completely eliminating a phylogroup becomes impossible because of the constant influx from the reservoir.

Using discrete-time simulations of the model with this colonization rate, we investigated how this external source impacted coexistence between two strains, by varying the proportion $q$ of clones provided by the reservoir between 0 (no reservoir) to 1 (only the reservoir). We consider here the same case

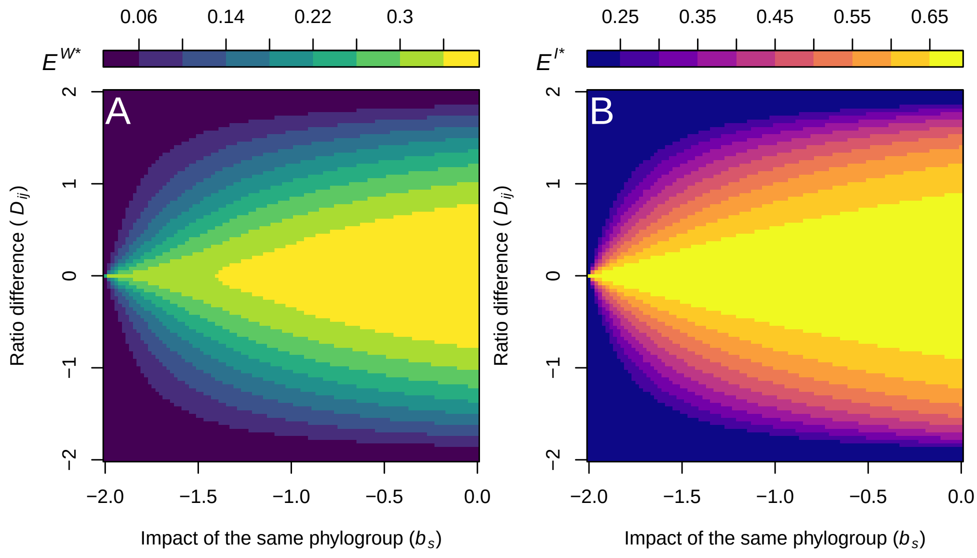


Figure S7: Asymptotic within host $E^{W*}$ (A) and inter-hosts $E^{I*}$ (B) coexistence as a function of the ratio difference $D_{i,j}$ and the proportion of clones provided by the reservoir $q$. For these simulations, $\overline{R_{i,j}} = 2$, $b = 0$, $d = 0$, $E^{W*} \in\ ]0,1]$ and $E^{I*} \in\ ]0, ln(2)]$.

scenario as the one presented in the main text of the article (see the Results section): a couple of phylogroups $i$ and $j$ characterized by an average colonization-to-clearance ratio $\overline{R_{i,j}} = (R_i + R_j)/2$ and a ratio difference $D_{i,j} = R_i - R_j$. We considered both phylogroups present at the same frequency in the reservoir $F_i^r = F_j^r = 1/2$, and computed the asymptotic within-host coexistence $E^{W*}$ and the asymptotic inter-host coexistence $E^{I*}$, with the same definition as presented in the main text of the article.

Results show that the reservoir facilitated coexistence (Fig S7). The range of values of $D_{i,j}$ values for which coexistence was stable widened dramatically as the proportion of colonization depending on the reservoir $q$ increased. This was the case both for the within-host coexistence $E^{W*}$ and the inter-host coexistence $E^{I*}$. These results show that even a reservoir with a somewhat limited impact on colonization would enable the coexistence of phylogroups with substantial differences in their colonization-to-clearance ratios. However, this effect would be mitigated if the fitter phylogroup were also the most abundant in the reservoir, e.g. if $R_i > R_j$ and $F_i^r > F_j^r$.

### 8. Supplementary figures

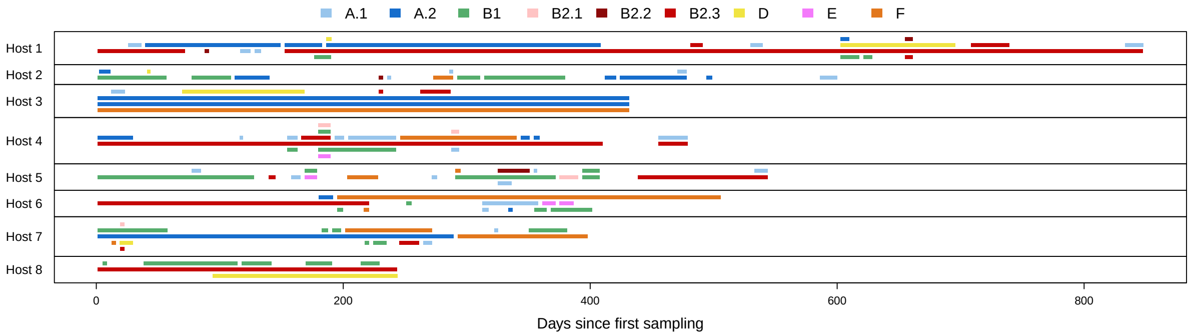


Figure S8: Presence of *E. coli* clones over time in each of the 8 hosts in the USA dataset. Each bar represents a single clone as defined in the study, starting either at 0, or midway between the first sampling including the clone and the previous one, and ending either at the date of the last sampling, or midway between the last sampling including the clone and the next one. Non-overlapping presence of several clones is represented on the same line for the sake of compactness, with no specific link between them.

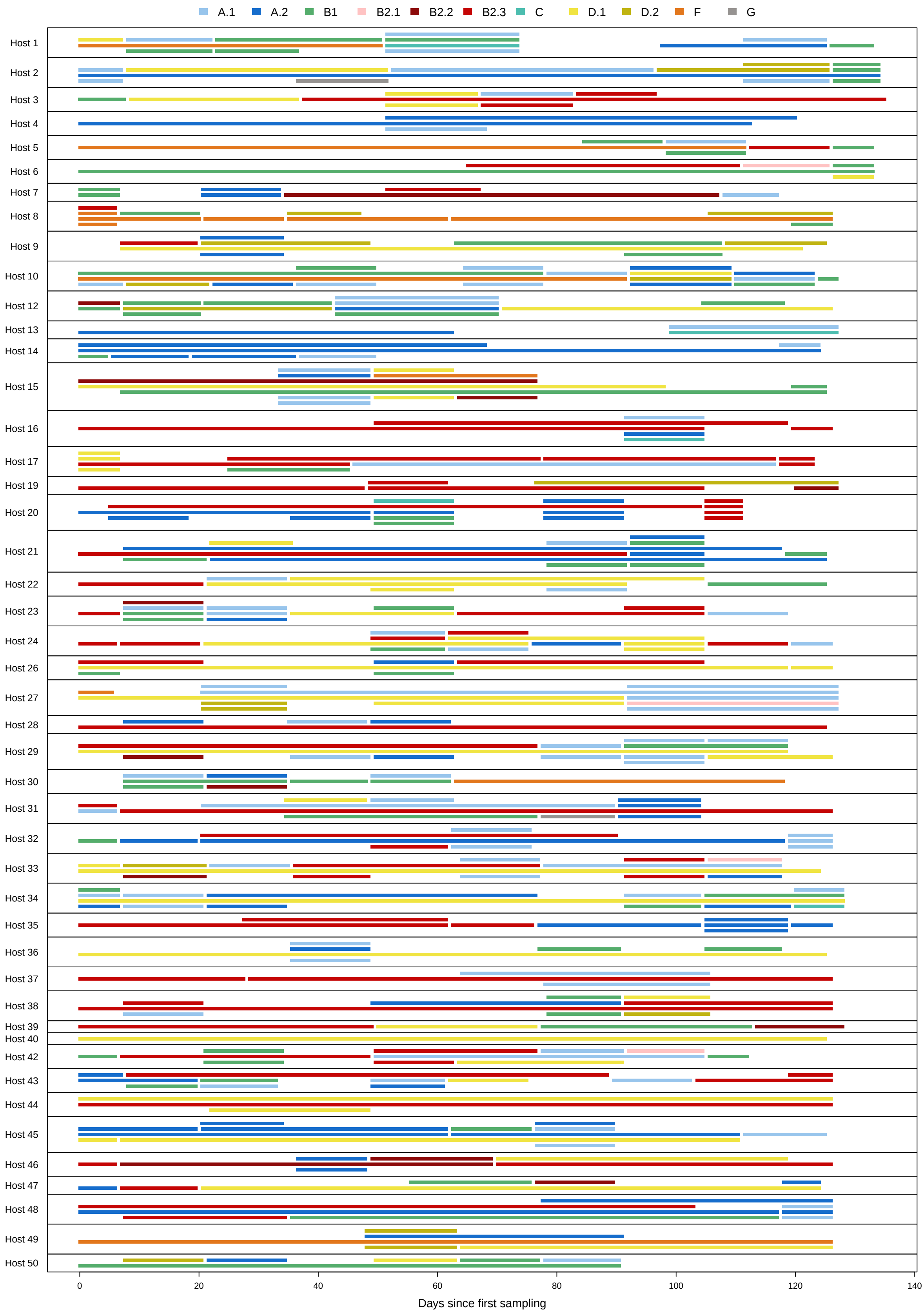


Figure S9: Presence of *E. coli* clones over time in each of the 46 hosts in the French dataset. Each bar represents a single clone as defined in the study, starting either at 0, or midway between the first sample including the clone and the previous one, and ending either at the date of the last sample, or midway between the last sample including the clone and the next one. Non-overlapping presence of several clones is represented on the same line for the sake of compactness, with no specific link between them.

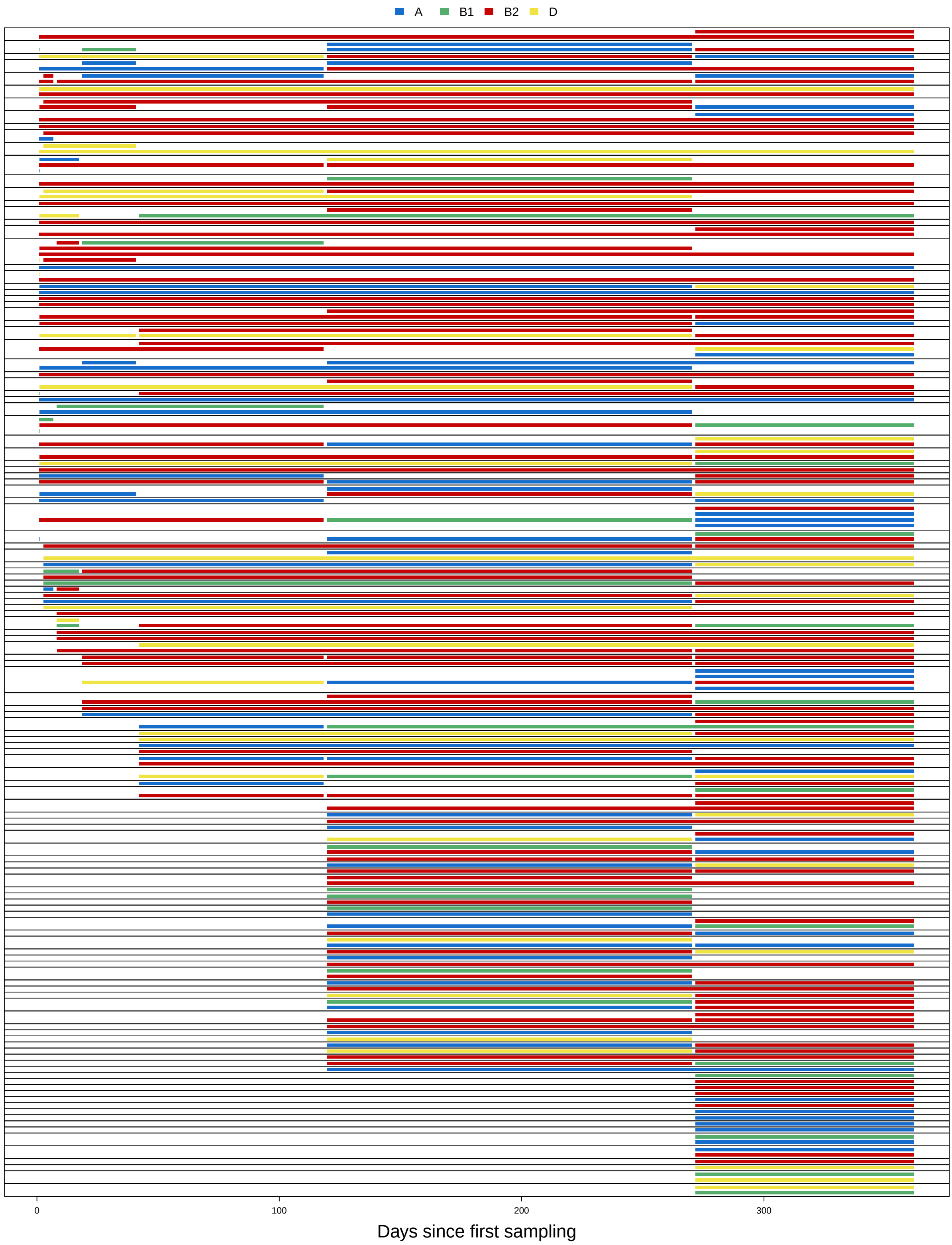


Figure S10: Presence of *E. coli* clones over time in each host of the infant (Sweden) dataset. Each bar represents a single clone as defined in the study, starting either at 0, or midway between the first sample including the clone and the previous one, and ending either at the date of the last sample, or midway between the last sample including the clone and the next one. Non-overlapping presence of several clones is represented on the same line for compactness, with no specific link between them. Hosts are separated by black lines.

### 9. Supplementary tables

#### 9.1. *AFT models for residence times*

##### 9.1.1 French dataset

| Model | Distribution | AIC | ΔAIC |
|---|---|---|---|
| $Y \sim hst + grp + ncl + ngrp$ | log-logistic | 622.501 | 0.000 |
| $Y \sim hst + grp + ncl + ngrp + log(dns)$ | log-logistic | 623.848 | 1.346 |
| $Y \sim hst + grp + ncl$ | log-logistic | 627.530 | 5.028 |
| $Y \sim hst + grp + ncl + log(dns)$ | log-logistic | 628.956 | 6.454 |

Table S1: AIC and ΔAIC of AFT models of residence for the French dataset, with ΔAIC<10, with either an exponential, Weibull or log-logistic distribution. The factors included are noted as follows: *hst* (host), *grp* (phylogroup), *ncl* (number of clones), *ngrp* (number of clones of the same phylogroup) and *dns* (total density).

| Coefficient | z-score | p-value |
|---|---|---|
| intercept (host 03/group D.2) | 7.578 | < 0.0001 |
| host 01 | 0.073 | 0.9415 |
| host 02 | 1.427 | 0.1535 |
| host 04 | 1.754 | 0.0795 |
| host 05 | -1.960 | 0.0500 |
| host 06 | -0.182 | 0.8559 |
| host 07 | 0.273 | 0.7852 |
| host 08 | -1.619 | 0.1054 |
| host 09 | 1.362 | 0.1733 |
| host 10 | -1.802 | 0.0716 |
| host 12 | 0.290 | 0.7717 |
| host 14 | -1.059 | 0.2897 |
| host 15 | -0.185 | 0.8533 |
| host 16 | 0.348 | 0.7276 |
| host 17 | 0.822 | 0.4110 |
| host 19 | 1.008 | 0.3134 |
| host 20 | 0.386 | 0.6992 |
| host 21 | 1.546 | 0.1222 |
| host 22 | 0.714 | 0.4750 |
| host 23 | -0.583 | 0.5597 |
| host 24 | -1.171 | 0.2416 |
| host 26 | -0.149 | 0.8812 |
| host 27 | 0.040 | 0.9684 |
| host 28 | -1.774 | 0.0760 |
| host 29 | -0.200 | 0.8413 |
| host 30 | -1.151 | 0.2497 |
| host 31 | 2.556 | 0.0106 |
| host 32 | 1.712 | 0.0869 |
| host 33 | -1.125 | 0.2606 |
| host 34 | 0.322 | 0.7477 |
| host 35 | -1.000 | 0.3174 |
| host 36 | -1.015 | 0.3101 |
| host 37 | 2.701 | 0.0069 |
| host 38 | -1.020 | 0.3079 |
| host 39 | -1.041 | 0.2979 |
| host 42 | -0.553 | 0.5799 |
| host 43 | -0.561 | 0.5747 |
| host 44 | 0.357 | 0.7210 |

| Coefficient | z-score | p-value |
|---|---|---|
| host 45 | 1.744 | 0.0812 |
| host 46 | 0.806 | 0.4200 |
| host 47 | 0.616 | 0.5378 |
| host 48 | 2.868 | 0.0041 |
| host 49 | 0.217 | 0.8282 |
| host 50 | -2.217 | 0.0267 |
| group A.1 | -2.311 | 0.0208 |
| group A.2 | 0.825 | 0.4094 |
| group B1 | -1.886 | 0.0593 |
| group B2.1 | -1.981 | 0.0476 |
| group B2.2 | 0.496 | 0.6201 |
| group B2.3 | 2.893 | 0.0038 |
| group C | -0.930 | 0.3522 |
| group D. | 3.303 | 0.0010 |
| group E | -1.437 | 0.1507 |
| group F | 3.230 | 0.0012 |
| group G | -3.336 | 0.0008 |
| number of clones | -4.218 | $< 0.0001$ |
| number of clones of the group | -2.654 | 0.0079 |
| Log. of scale | -10.912 | $< 0.0001$ |

Table S2: Coefficients of the selected AFT model of residence for the French dataset, with their associated z-scores and p-values. The model intercept uses host 03 and phylogroup D.2 as reference, as they are the closest to the average estimate among hosts and among groups, respectively. The number of clones and number of clones of the group were scaled prior to running the model, by subtracting their mean value and dividing them by their standard deviation.

##### 9.1.2 USA dataset

| Model | Distribution | AIC | ΔAIC |
|---|---|---|---|
| $Y \sim hst + grp + ncl + ngrp + log(dscl)$ | exponential | 469.141 | 0.000 |
| $Y \sim hst + grp + ncl + ngrp + log(dns) + log(dscl)$ | exponential | 470.128 | 0.987 |
| $Y \sim hst + grp + ncl + ngrp + log(dscl)$ | Weibull | 470.555 | 1.414 |
| $Y \sim hst + grp + ncl + ngrp$ | exponential | 471.031 | 1.891 |
| $Y \sim hst + grp + ncl + log(dscl)$ | exponential | 471.392 | 2.252 |
| $Y \sim hst + grp + ncl + ngrp + log(dns) + log(dscl)$ | Weibull | 471.402 | 2.261 |
| $Y \sim hst + grp + ncl + ngrp + log(dns)$ | exponential | 471.890 | 2.749 |
| $Y \sim hst + grp + ncl + log(dscl)$ | Weibull | 472.579 | 3.438 |
| $Y \sim hst + grp + ncl + ngrp$ | Weibull | 472.610 | 3.469 |
| $Y \sim hst + grp + ncl + log(dns) + log(dscl)$ | exponential | 472.991 | 3.850 |
| $Y \sim hst + grp + ncl + ngrp + log(dns) + log(dscl)$ | log-logistic | 472.998 | 3.857 |
| $Y \sim hst + grp + ncl$ | exponential | 473.381 | 4.240 |
| $Y \sim hst + grp + ncl + ngrp + log(dns)$ | Weibull | 473.468 | 4.327 |
| $Y \sim hst + grp + ncl + log(dns)$ | exponential | 473.742 | 4.601 |
| $Y \sim hst + grp + ncl + log(dns) + log(dscl)$ | Weibull | 474.047 | 4.906 |
| $Y \sim hst + grp + ncl$ | Weibull | 474.763 | 5.622 |
| $Y \sim hst + grp + ncl + log(dns)$ | Weibull | 475.144 | 6.003 |
| $Y \sim hst + grp + ngrp + log(dns) + log(dscl)$ | exponential | 475.178 | 6.037 |
| $Y \sim hst + grp + ncl + log(dns) + log(dscl)$ | log-logistic | 475.351 | 6.210 |
| $Y \sim hst + grp + ncl + ngrp + log(dscl)$ | log-logistic | 475.829 | 6.688 |
| $Y \sim hst + grp + ngrp + log(dns) + log(dscl)$ | Weibull | 475.990 | 6.849 |
| $Y \sim hst + grp + ncl + log(dscl)$ | log-logistic | 477.349 | 8.209 |

Table S3: AIC and ΔAIC of AFT models of residence for the USA dataset, with ΔAIC<10, with either an exponential, Weibull or log-logistic distribution. The factors included are noted as follows: *hst* (host), *grp* (phylogroup), *ncl* (number of clones), *ngrp* (number of clones of the same phylogroup), *dns* (total density) and *dscl* (total density of the clone).

| Coefficient | z-score | p-value |
|---|---|---|
| intercept (host 5/group A.2) | 8.661 | $< 0.0001$ |
| host 1 | 3.007 | 0.0026 |
| host 2 | -0.926 | 0.3546 |
| host 3 | -0.141 | 0.8875 |
| host 4 | -1.985 | 0.0471 |
| host 6 | 1.311 | 0.1898 |
| host 7 | -3.822 | 0.0001 |
| host 8 | 1.810 | 0.0702 |
| group A.1 | -3.496 | 0.0005 |
| group B1 | -0.990 | 0.3224 |
| group B2.1 | -1.136 | 0.2558 |
| group B2.2 | -3.103 | 0.0019 |
| group B2.3 | 4.916 | $< 0.0001$ |
| group D | 1.618 | 0.1056 |
| group E | -2.044 | 0.0409 |
| group F | 3.679 | 0.0002 |
| number of clones | -3.444 | 0.0006 |
| number of clones of the group | -2.135 | 0.0328 |
| log. of clone density | 1.983 | 0.0473 |

Table S4: Coefficients of the selected AFT model of residence for the USA dataset, with their associated z-scores and p-values. The model intercept uses host 5 and phylogroup A.2 as reference, as they are the closest to the average estimate among hosts and among groups, respectively. The number of clones and number of clones of the group were scaled prior to running the model, by subtracting their mean value and dividing them by their standard deviation.

##### 9.1.3 Infant dataset

| Model | Distribution | AIC | ΔAIC |
|---|---|---|---|
| $Y \sim grp + ncl$ | Weibull | 303.041 | 0.000 |
| $Y \sim grp + ncl + log(dns)$ | Weibull | 304.851 | 1.810 |
| $Y \sim grp + ncl + log(dscl)$ | Weibull | 304.907 | 1.865 |
| $Y \sim grp + ncl + ngrp$ | Weibull | 304.976 | 1.935 |
| $Y \sim grp + ngrp$ | Weibull | 305.398 | 2.357 |
| $Y \sim grp$ | Weibull | 305.743 | 2.701 |
| $Y \sim grp + ncl + ngrp + log(dns)$ | Weibull | 306.804 | 3.763 |
| $Y \sim grp + ncl + log(dns) + log(dscl)$ | Weibull | 306.843 | 3.802 |
| $Y \sim grp + ncl + ngrp + log(dscl)$ | Weibull | 306.860 | 3.818 |
| $Y \sim grp + ngrp + log(dns)$ | Weibull | 307.323 | 4.282 |
| $Y \sim grp + ngrp + log(dscl)$ | Weibull | 307.366 | 4.324 |
| $Y \sim grp + log(dns) + log(dscl)$ | Weibull | 307.556 | 4.515 |
| $Y \sim grp + log(dscl)$ | Weibull | 307.607 | 4.566 |
| $Y \sim grp + log(dns)$ | Weibull | 307.639 | 4.598 |
| $Y \sim grp + ngrp + log(dns) + log(dscl)$ | Weibull | 308.373 | 5.331 |
| $Y \sim grp + ncl + ngrp + log(dns) + log(dscl)$ | Weibull | 308.793 | 5.752 |

Table S5: AIC and ΔAIC of AFT models of residence for the infant dataset, with ΔAIC<10, with either an exponential, Weibull or log-logistic distribution. The factors included are noted as follows: *hst* (host), *grp* (phylogroup), *ncl* (number of clones), *ngrp* (number of clones of the same phylogroup), *dns* (total density) and *dscl* (total density of the clone).

#### 9.2. *AFT models for colonization rates*

##### 9.2.1 French dataset

| Model | Distribution | AIC | ΔAIC |
|---|---|---|---|
| $Y \sim hst + grp + ncl$ | Weibull | 436.443 | 0.000 |
| $Y \sim hst + grp + ncl + ngrp$ | Weibull | 437.722 | 1.279 |
| $Y \sim hst + grp + ncl + log(dns)$ | Weibull | 437.987 | 1.544 |
| $Y \sim hst + grp + ncl + ngrp + log(dns)$ | Weibull | 439.252 | 2.809 |
| $Y \sim hst + grp + ncl$ | log-logistic | 440.191 | 3.748 |
| $Y \sim hst + grp + ncl + ngrp$ | log-logistic | 441.242 | 4.798 |
| $Y \sim hst + grp + ncl + log(dns)$ | log-logistic | 441.395 | 4.952 |
| $Y \sim hst + grp + ncl + ngrp + log(dns)$ | log-logistic | 442.435 | 5.992 |

Table S6: AIC and ΔAIC of AFT models of colonization for the French dataset, with ΔAIC<10, with either an exponential, Weibull or log-logistic distribution. The factors included are noted as follows: *hst* (host), *grp* (phylogroup), *ncl* (number of clones), *ngrp* (number of clones of the same phylogroup) and *dns* (total density).

| Coefficient | z-score | p-value |
|---|---|---|
| intercept (host 21/group D.2) | 9.157 | < 0.0001 |
| host 01 | -0.010 | 0.9916 |
| host 02 | 0.689 | 0.4906 |
| host 03 | -0.915 | 0.3600 |
| host 05 | -1.558 | 0.1192 |
| host 06 | -0.761 | 0.4465 |
| host 07 | 0.548 | 0.5834 |
| host 08 | 0.713 | 0.4760 |
| host 09 | -0.099 | 0.9215 |
| host 10 | -1.584 | 0.1132 |
| host 12 | -0.535 | 0.5924 |
| host 14 | 1.302 | 0.1930 |
| host 15 | 0.367 | 0.7138 |
| host 16 | 1.027 | 0.3043 |
| host 17 | 1.945 | 0.0518 |
| host 19 | 0.782 | 0.4344 |
| host 20 | -0.487 | 0.6260 |
| host 22 | -0.166 | 0.8684 |
| host 23 | -1.212 | 0.2255 |
| host 24 | -1.029 | 0.3034 |
| host 26 | 2.978 | 0.0029 |
| host 27 | 0.265 | 0.7909 |
| host 28 | -0.803 | 0.4217 |
| host 29 | -0.895 | 0.3707 |
| host 30 | -2.247 | 0.0246 |
| host 31 | -1.388 | 0.1650 |
| host 32 | 1.183 | 0.2369 |
| host 33 | -1.704 | 0.0884 |
| host 34 | 1.903 | 0.0571 |
| host 35 | -0.056 | 0.9557 |
| host 36 | 0.162 | 0.8710 |
| host 37 | -0.828 | 0.4076 |
| host 38 | 0.275 | 0.7836 |
| host 39 | -0.107 | 0.9144 |
| host 42 | -1.458 | 0.1448 |
| host 43 | -0.981 | 0.3264 |
| host 45 | 2.884 | 0.0039 |
| host 46 | 0.019 | 0.9845 |

| Coefficient | z-score | p-value |
|---|---|---|
| host 47 | -0.087 | 0.9310 |
| host 48 | 2.605 | 0.0092 |
| host 49 | 0.308 | 0.7581 |
| host 50 | -0.698 | 0.4850 |
| group A.1 | 1.309 | 0.1904 |
| group A.2 | 6.781 | $< 0.0001$ |
| group B1 | 5.493 | $< 0.0001$ |
| group B2.1 | -4.931 | $< 0.0001$ |
| group B2.2 | -2.983 | 0.0029 |
| group B2.3 | 9.335 | $< 0.0001$ |
| group C | -3.030 | 0.0024 |
| group D.1 | 1.492 | 0.1356 |
| group E | -2.771 | 0.0056 |
| group F | 1.240 | 0.2151 |
| group G | -2.371 | 0.0177 |
| number of clones | -9.493 | $< 0.0001$ |
| Log. of scale | -12.918 | $< 0.0001$ |

Table S7: Coefficients of the selected AFT model of colonization for the French dataset, with their associated z-scores and p-values. The model intercept uses host 21 and phylogroup D.2 as reference, as they are the closest to the average estimate among hosts and among groups, respectively. The number of clones and number of clones of the group were scaled prior to running the model, by subtracting their mean value and dividing them by their standard deviation.

##### 9.2.2 USA dataset

| Model | Distribution | AIC | ΔAIC |
|---|---|---|---|
| $Y \sim hst + grp + ncl$ | Weibull | 481.535 | 0.000 |
| $Y \sim hst + grp + ncl + ngrp$ | Weibull | 481.720 | 0.185 |
| $Y \sim hst + grp + ncl + log(dns)$ | Weibull | 482.886 | 1.350 |
| $Y \sim hst + grp + ncl + ngrp + log(dns)$ | Weibull | 483.002 | 1.466 |
| $Y \sim hst + grp + ncl + ngrp$ | log-logistic | 489.479 | 7.944 |
| $Y \sim hst + grp + ncl$ | log-logistic | 489.599 | 8.064 |
| $Y \sim hst + grp + ncl + ngrp + log(dns)$ | log-logistic | 491.254 | 9.719 |
| $Y \sim hst + grp + ncl + log(dns)$ | log-logistic | 491.473 | 9.938 |

Table S8: AIC and ΔAIC of AFT models of colonization for the USA dataset, with ΔAIC<10, with either an exponential, Weibull or log-logistic distribution. The factors included are noted as follows: *hst* (host), *grp* (phylogroup), *ncl* (number of clones), *ngrp* (number of clones of the same phylogroup), *dns* (total density) and *dscl* (total density of the clone).

| Coefficient | z-score | p-value |
|---|---|---|
| intercept (host 6/group B1) | 17.879 | $< 0.0001$ |
| host 1 | 1.698 | 0.0896 |
| host 2 | -1.475 | 0.1403 |
| host 3 | 2.950 | 0.0032 |
| host 4 | -1.451 | 0.1469 |
| host 5 | -2.826 | 0.0047 |
| host 7 | 2.445 | 0.0145 |
| host 8 | 2.612 | 0.0090 |
| group A.1 | -6.219 | $< 0.0001$ |
| group A.2 | 3.874 | 0.0001 |
| group B2.1 | -10.459 | $< 0.0001$ |
| group B2.2 | -7.768 | $< 0.0001$ |
| group B2.3 | 8.472 | $< 0.0001$ |
| group D | 1.686 | 0.0918 |
| group E | -6.837 | $< 0.0001$ |
| group F | -0.824 | 0.4099 |
| number of clones | -9.775 | $< 0.0001$ |
| Log. of scale | -4.987 | $< 0.0001$ |

Table S9: Coefficients of the selected AFT model of colonization for the USA dataset, with their associated z-scores and p-values. The model intercept uses host 6 and phylogroup B1 as reference, as they are the closest to the average estimate among hosts and among groups, respectively. The number of clones and number of clones of the group were scaled prior to running the model, by subtracting their mean value and dividing them by their standard deviation.

##### 9.2.3 Infant dataset

| Model | Distribution | AIC | ΔAIC |
|---|---|---|---|
| $Y \sim grp + ncl$ | Weibull | 205.132 | 0.000 |
| $Y \sim grp + ncl + log(dns)$ | Weibull | 206.868 | 1.737 |
| $Y \sim grp + ncl + ngrp$ | Weibull | 207.047 | 1.915 |
| $Y \sim grp$ | Weibull | 207.242 | 2.110 |
| $Y \sim grp + ncl + ngrp + log(dns)$ | Weibull | 208.687 | 3.556 |
| $Y \sim grp + log(dns)$ | Weibull | 208.805 | 3.674 |
| $Y \sim grp + ngrp$ | Weibull | 208.942 | 3.810 |
| $Y \sim grp + ngrp + log(dns)$ | Weibull | 210.304 | 5.172 |

Table S10: AIC and ΔAIC of AFT models of colonization for the USA dataset, with ΔAIC<10, with either an exponential, Weibull or log-logistic distribution. The factors included are noted as follows: *hst* (host), *grp* (phylogroup), *ncl* (number of clones), *ngrp* (number of clones of the same phylogroup) and *dns* (total density).

#### 9.3. *Cox models for residence times*

##### 9.3.1 French dataset

| Coefficient | z-score | p-value |
|---|---|---|
| group A.1 | 0.326 | 0.7442 |
| group A.2 | -0.609 | 0.5424 |
| group B1 | 0.904 | 0.3662 |
| group B2.1 | 1.587 | 0.1126 |
| group B2.2 | -0.543 | 0.5874 |
| group B2.3 | -2.685 | 0.0073 |
| group C | 0.138 | 0.8905 |
| group D.1 | -3.108 | 0.0019 |
| group E | 0.967 | 0.3333 |
| group F | -1.761 | 0.0783 |
| group G | 1.859 | 0.0630 |
| number of clones | 2.463 | 0.0143 |
| number of clones of the group | 3.431 | 0.0006 |

Table S11: Coefficients of the Cox model of residence for the USA dataset, with their associated z-scores and p-values. The model is stratified by host, as this factor did not respect the proportional hazard hypothesis. The model intercept uses phylogroup D.2 as the corresponding AFT model. The number of clones and number of clones of the group were scaled prior to running the model, by subtracting their mean value and dividing them by their standard deviation.

##### 9.3.2 USA dataset

| Coefficient | z-score | p-value |
|---|---|---|
| group A.1 | 2.214 | 0.0268 |
| group B1 | 0.192 | 0.8476 |
| group B2.1 | 1.045 | 0.2961 |
| group B2.2 | 1.648 | 0.0994 |
| group B2.3 | -2.535 | 0.0113 |
| group D | -1.624 | 0.1044 |
| group E | 0.955 | 0.3397 |
| group F | -3.093 | 0.0020 |
| number of clones | 3.457 | 0.0005 |
| number of clones of the group | 2.068 | 0.0285 |
| log. of clone density | -2.271 | 0.0231 |

Table S12: Coefficients of the Cox model of residence for the USA dataset, with their associated z-scores and p-values. The model is stratified by host, as this factor did not respect the proportional hazard hypothesis. The model intercept uses phylogroup A.2 as the corresponding AFT model. The number of clones and number of clones of the group were scaled prior to running the model, by subtracting their mean value and dividing them by their standard deviation.

#### 9.4. *Cox models for colonization rates*

##### 9.4.1 French dataset

| Coefficient | z-score | p-value |
|---|---|---|
| group A.1 | -0.320 | 0.7492 |
| group A.2 | -4.056 | $< 0.0001$ |
| group B1 | -2.421 | 0.0155 |
| group B2.1 | 0.002 | 0.9985 |
| group B2.2 | 0.005 | 0.9961 |
| group B2.3 | -5.708 | $< 0.0001$ |
| group C | 0.001 | 0.9996 |
| group D.1 | -0.540 | 0.5889 |
| group E | 2.619 | 0.0088 |
| group F | -2.655 | 0.0079 |
| group G | 0.001 | 0.9996 |
| number of clones | 5.403 | $< 0.0001$ |

Table S13: Coefficients of Cox model of colonization rate for the French dataset, with their associated z-scores and p-values. The model is stratified by host, as this factor did not respect the proportional hazard hypothesis. The model intercept uses phylogroup D.2 as the corresponding AFT model. The number of clones and number of clones of the group were scaled prior to running the model, by subtracting their mean value and dividing them by their standard deviation.

##### 9.4.2 USA dataset

| Coefficient | z-score | p-value |
|---|---|---|
| group A.1 | 3.511 | 0.0002 |
| group B1 | -2.338 | 0.0097 |
| group B2.1 | 6.794 | $< 0.0001$ |
| group B2.2 | 3.476 | 0.0003 |
| group B2.3 | -3.498 | 0.0002 |
| group D | -1.211 | 0.1130 |
| group E | 6.687 | $< 0.0001$ |
| group F | -0.811 | 0.2086 |
| number of clones | 3.326 | 0.0004 |

Table S14: Coefficients of the Cox model of colonization for the USA dataset, with their associated z-scores and p-values. The model is stratified by host, as this factor did not respect the proportional hazard hypothesis. The model intercept uses phylogroup A.2 as the corresponding AFT model. The number of clones and number of clones of the group were scaled prior to running the model, by subtracting their mean value and dividing them by their standard deviation.
